# Generation and application of human pluripotent stem cells derived peri-nervoids

**DOI:** 10.64898/2026.09.14.750880

**Authors:** Yanjun Guan, Ruichao He, Ziyue Xiang, Yiben Ouyang, Haolin Liu, Shengfang Zhang, Xinyao Liu, Chang Liu, Zhixin Wang, Yu Wang, Shanlin Chen

## Abstract

Autologous nerve grafts remain the gold standard for peripheral nerve repair, but donor scarcity, donor-site morbidity and incomplete functional recovery limit their use. Schwann cells (SCs) drive autograft-mediated regeneration, yet their limited availability and expansion impede SC-based therapies. Here, we generated human pluripotent stem cell-derived SCs, motor neuron-enriched spinal cord organoids and nociceptive sensory neuron-enriched dorsal root ganglion organoids, and assembled them into a nerve-like construct termed the Peri-nervoid. Single-cell RNA sequencing revealed context-dependent SC plasticity: axonal co-culture promoted myelination, whereas axonal transection induced a repair phenotype resembling SC dedifferentiation during Wallerian degeneration. Following transplantation into 7-mm sciatic nerve defects in NSG mice, human SCs survived and remyelinated regenerating host axons. No abnormal proliferation, systemic toxicity or major-organ histopathology was detected. These findings position the Peri-nervoid as a scalable, developmentally inspired platform for developing engineered grafts for peripheral nerve repair.

## Introduction

Autologous nerve transplantation is widely accepted in clinical practice for peripheral nerve defects exceeding 3cm. The superiority of autografts lies not only in their native structure, but in the living cellular microenvironment they provide. Specifically, autografts contain viable Schwann cells (SCs) within intact basal lamina tubes^1^. Following transplantation and loss of axonal contact, these SCs rapidly reprogram into a repair phenotype: they clear myelin debris, upregulate neurotrophic factors, and organize into longitudinal Bands of Büngner that guide regenerating axons across the graft^2–5^. This transition is a central cellular response during Wallerian degeneration^6,7^. The structural guidance and adaptive cellular support provided by autografts together account for their regenerative efficacy. However, limited donor nerve availability, donor-site morbidity, and mismatches in calibre and fascicular architecture between donor and recipient nerves restrict their broader clinical use^8,9^. Developing alternatives that reproduce both the structural and cellular functions of autografts has therefore remained a central goal in peripheral nerve repair.

Strategies incorporating exogenous SCs transplantation have been pursued for decades. SCs isolated from adult human peripheral nerves can survive after transplantation into rodent sciatic nerve defects and contribute to the myelination of regenerating axons^10,11^. However, clinical translation remains challenging because sufficient numbers of SCs must be obtained while preserving their identity and regenerative function during isolation and ex vivo expansion^12^. Moreover, many experimental approaches deliver SCs as cell suspensions. Although this method facilitates cell distribution within the lesion, SCs alone do not provide the aligned structural support required to bridge a segmental nerve defect and guide regenerating axons. Existing approaches therefore often reproduce only part of the regenerative environment of an autograft: acellular conduits provide physical guidance but lack injury-responsive cells, whereas SC suspensions provide cellular support without preformed axonal architecture. An engineered graft combining organised structural guidance with responsive SCs may more closely reproduce the regenerative functions of an autologous nerve graft.

Human pluripotent stem cells (hPSCs) provide a potentially scalable source of SCs^13,14^. Recent studies have demonstrated that human induced pluripotent stem cells (hiPSC)-derived SCs closely resemble primary SCs in gene expression and functional properties, and can be expanded in vitro with stable phenotype, offering a practical solution to the cell sourcing problem^14,15^. With a reliable SC source established, we have turned our attention toward biomimicry of the native architecture of peripheral nerves. Bi-directional signaling between axons and SCs, mediated by factors such as axon-derived neuregulin and biophysical cues, regulates SCs survival, proliferation, and myelination^16,17^. Following the loss of axonal contact, SCs adopt a repair phenotype and form Bands of Büngner, but prolonged period without axons progressively reduces their regenerative capacity^18^. Hence, we reasoned that combining SCs with neuronal axons could reproduce key structural features of peripheral nerves while supporting SC function through reciprocal axon-SC interactions. In parallel, protocols for generating hPSC-derived neural organoid have developed considerably^19,20^. Spinal cord (motor neuron) and dorsal root ganglion (sensory neuron) organoids can extend centimeter-scale neurite bundles in vitro^21,22^, providing suitable axonal components for constructing a nerve substitute.

Peripheral nerve development provides another rationale for this approach. Sensory axons from dorsal root ganglion neurons and motor axons from spinal cord neurons extend towards peripheral targets, while neural crest-derived Schwann cell precursors migrate along nascent axons, differentiate and organize myelinated and unmyelinated fibers. This coordinated developmental process suggests that combining hPSC-derived SCs with motor and sensory axons may generate a preformed nerve-like tissue in which reciprocal axon-SC interactions support tissue organization and regenerative function.

Here, we developed the Peri-nervoid, a preformed nerve-like graft composed of hPSC-derived SCs and axonal bundles extending from motor neuron-enriched spinal cord organoids (SCOs) and nociceptive sensory neuron-enriched dorsal root ganglion organoids (DRGOs). We used single-cell RNA sequencing to determine how motor axons, sensory axons and axonal injury influence SC states, and evaluated graft function and safety in a 7-mm sciatic nerve defect in NSG mice. By integrating expandable human SCs with organised axonal architecture, the Peri-nervoid provides a model for investigating axon-SC interactions and a candidate graft for peripheral nerve repair.

## Results

### 1. Establishment of a three-component hiPSC-derived neural platform for peri-nervoid assembly

To generate the cellular building blocks for Peri-nervoid assembly, we first established protocols to derive three major neural lineages from both Embryonic Stem Cells (ESCs) and human induced pluripotent stem cells (hiPSCs): spinal cord organoids (SCOs) enriched for motor neurons, dorsal root ganglion organoids (DRGOs) enriched for sensory neurons, and purified Schwann cells (SCs). For SCO generation, embryoid bodies (EBs) were sequentially directed toward neuromesodermal progenitors (NMPs), neural progenitors (NPs), and ventral spinal motor neurons (MNs) through timed exposure to small molecules and growth factors (Fig. 1A). Bright-field imaging showed progressive morphological maturation from regular spherical structures at D0 to prominent neuroepithelial-like structures by D20, with further size and complexity increases by D70 (Fig. 1B). Real-time PCR analysis demonstrated dynamic expression of stage-specific markers: neural ectoderm markers (*SOX2, PAX6*) peaked at D5 and declined thereafter, ventral motor neuron progenitor markers (*NKX6-1, OLIG2*) increased through D10–D20 (Fig. 1C). Immunofluorescence staining on D30 cryosections confirmed abundant TUJ1⁺neurons with widespread ISL1⁺motor neurons and FOXP1⁺/LIM1⁺subpopulations (Fig. 1D). Whole-mount 3D imaging of cleared D70 SCOs exhibited dense neuronal networks with peripheral somata and interwoven internal axon bundles, clearly showing ISL1⁺somata extending TUJ1⁺neurites (Fig. 1E). Functional maturation was confirmed by spontaneous calcium oscillations at D40 (Fig. 1F) and MEA recordings showing progressive increases in firing frequency, synchrony, and spike amplitude from D40 to D50 (Fig. 1G).

**Fig. 1.**
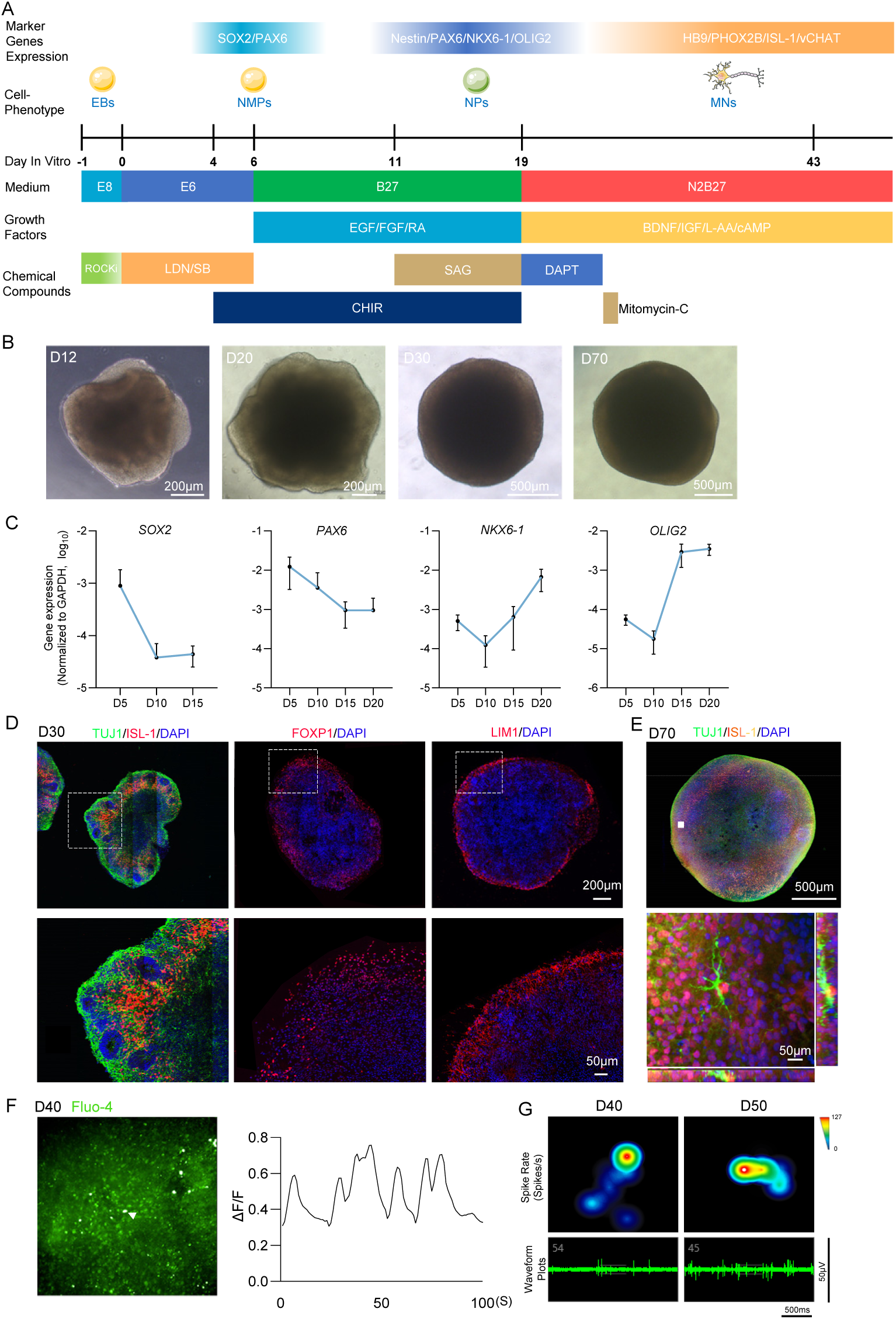
Generation and characterization of spinal cord organoids (SCOs) with motor neuron identity and functional electrical activity. **A.** Schematic illustration of the differentiation protocol for generating SCOs from hiPSCs. EB: embryoid body; NMP: neuromesodermal progenitor; NP: neural progenitor; MN: motor neuron; ROCKi: ROCK inhibitor. **B.** Bright-field images of SCOs during the differentiation process. **C.** Temporal expression profiles of motor neuron progenitor and terminal differentiation markers in SCOs during in vitro differentiation. Data are shown as mean ± SD from 3 independent differentiations (n = 3 biological replicates per time point; N= 9 total; one-way ANOVA: F (2, 23) = 7.31; P = 0.003 for *SOX2*; F (3, 30) = 9.06, P < 0.001 for *PAX6*; F (3, 30) = 22.4; P < 0.001 for *NKX6-1*, F (3, 30) = 26.6; P < 0.001 for *OLIG2*). **D.** Immunofluorescence staining in SCOs cryo-sections at day 30 of in vitro differentiation showing expression of motor neuron markers (TUJ1, ISL1, FOXP1, LIM1). **E.** Whole-mount 3D confocal imaging of an intact SCO immunolabeled for TUJ1 (neurons, green) and ISL-1 (motor neurons, red–yellow gradient) after 3DISCO clearing. Lower panels show higher-magnification views of the regions indicated by the white rectangles. **F.** Representative spontaneous calcium transients in SCOs at day 40. ΔF/F represents fluorescence intensity normalized to baseline; the white arrow indicates the cell analyzed. **G.** MEA recordings of SCOs at day 40 and day 50. Top: heatmaps of normalized firing rates; bottom: representative voltage spike waveforms.

For DRGO generation, EBs were similarly patterned toward neural crest (NC) cells and sensory neurons (SNs) (Fig. 2A). Bright-field imaging showed that DRGOs transitioned from regular spherical structures at D0 through phase-bright neuroepithelial-like structures by D10, with a noticeable clearance of non-neuronal cell debris between D5 and D10 (Fig. 2B). qPCR analysis confirmed the progressive differentiation of DRGOs from neural crest progenitors (*SOX10*) through sensory neuron progenitors (*NGN1/2*) to mature sensory neurons expressing *NRG1*, *GFRA2*, and *vGLUT2* (Fig. 2C). Immunofluorescence staining confirmed abundant TUJ1^+^ neurons co-expressing PRPH, SOX10, BRN3A, RUNX3 and PVALB, with BRN3A/RUNX3 positivity suggesting the presence of nociceptive and mechanosensitive neuronal subtypes within the DRGOs (Fig. 2D). Calcium imaging and MEA recordings confirmed spontaneous activity and progressively maturing electrical properties in DRGOs by D45–D50 (Fig. 2E–F).

**Fig. 2.**
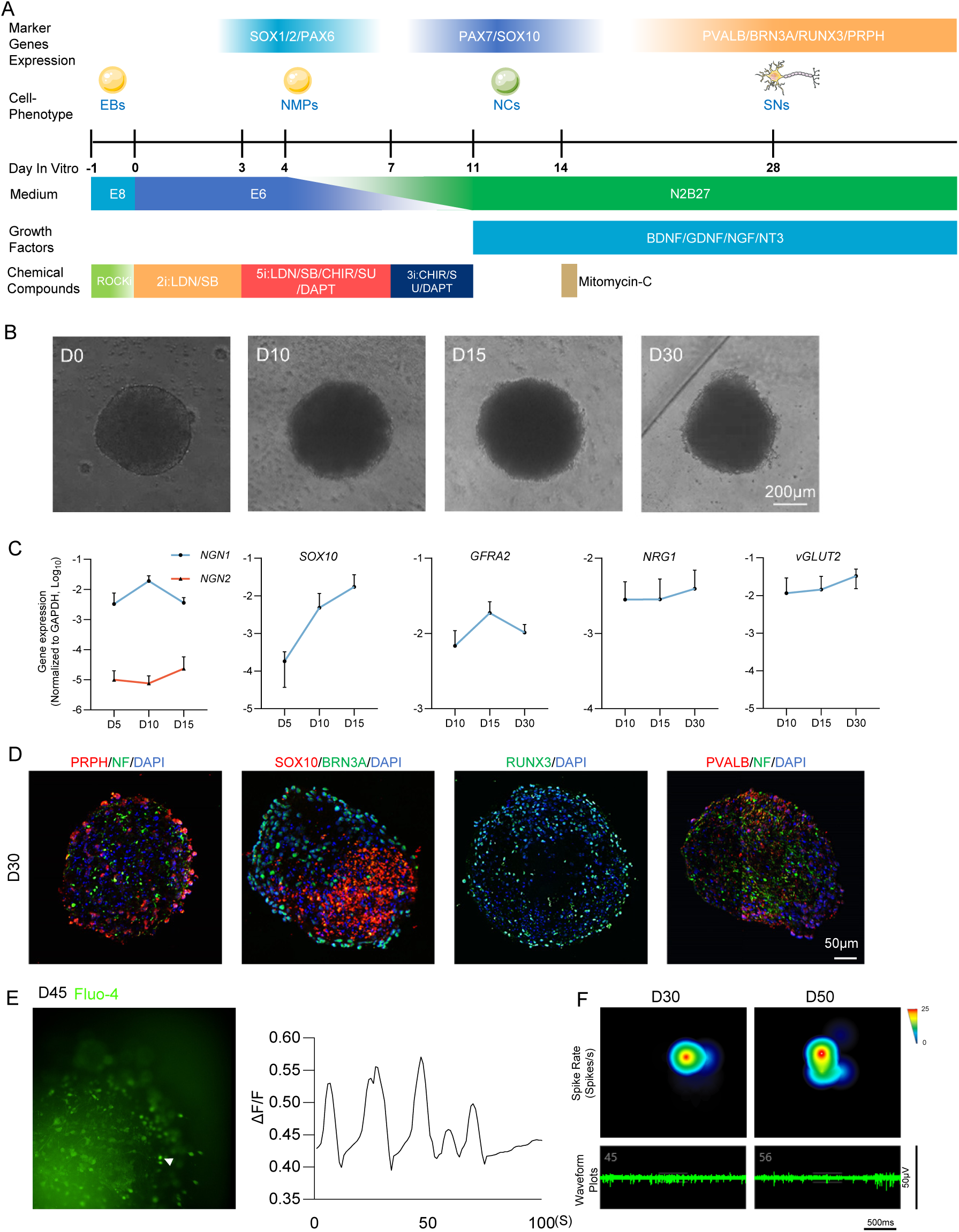
Generation and characterization of dorsal root ganglion organoids (DRGOs) with sensory neuron identity and functional electrical activity. A. Schematic illustration of the differentiation protocol for generating DRGOs from hiPSCs. NC: neural crest cells; SNs: sensory neurons. B. Brightfield images of DRGOs at indicated time points during differentiation. C. Temporal expression profiles of sensory neuron progenitor and terminal differentiation markers in DRGOs during in vitro differentiation. Data are shown as mean ± SD from 3 independent differentiations (n = 4 biological replicates per time point; N = 12 total; one-way ANOVA: F (2, 33) = 25.9; P < 0.001 for *NGN1*, F (2, 33) = 1.91; P = 0.165 for *NGN2*, F (2, 33) = 7.015; P = 0.0029 for *SOX10*, F (2, 33) = 16.02; P < 0.0001 for *GFRA2,* F (2, 33) = 0.759; P = 0.476 for *NRG1,* F (2, 33) = 5.11; P = 0.012 for *vGLUT2*). D. Immunofluorescence staining on DRGO cryosections at day 30 showing sensory neuron markers (PRPH, BRN3A, RUNX3, and PVALB). E. Representative spontaneous calcium transients in DRGOs at day 45. ΔF/F represents fluorescence intensity normalized to baseline. F. MEA recordings of DRGOs at day 30 and day 50. Top: heatmaps of normalized firing rates; bottom: representative voltage spike waveforms.

For SC derivation, monolayer cultures were directed through NC and Schwann cell progenitor (SCP) stages, followed by FACS purification and expansion (Fig. 3A). Bright-field imaging captured the morphological transition from epithelial-like colonies at D3 to bipolar/spindle-shaped progenitors at D10–D15, with homogeneous SC morphology after sorting (P1) that was stably maintained through P15 (Fig. 3B, F). qPCR showed sequential expression of NC/SCP markers (*PAX6, SOX10*), and mature SCs markers (*S100β*) (Fig. 3C). FACS confirmed that CD49d⁺ SCs purity around 10-30% across two independent lines (Fig. 3D). Immunofluorescence confirmed robust co-expression of SOX10, S100β, and NGFR in sorted cells (Fig. 3E), with stable maintenance of SOX10/S100β expression through long-term expansion (Fig. 3G).

**Fig. 3.**
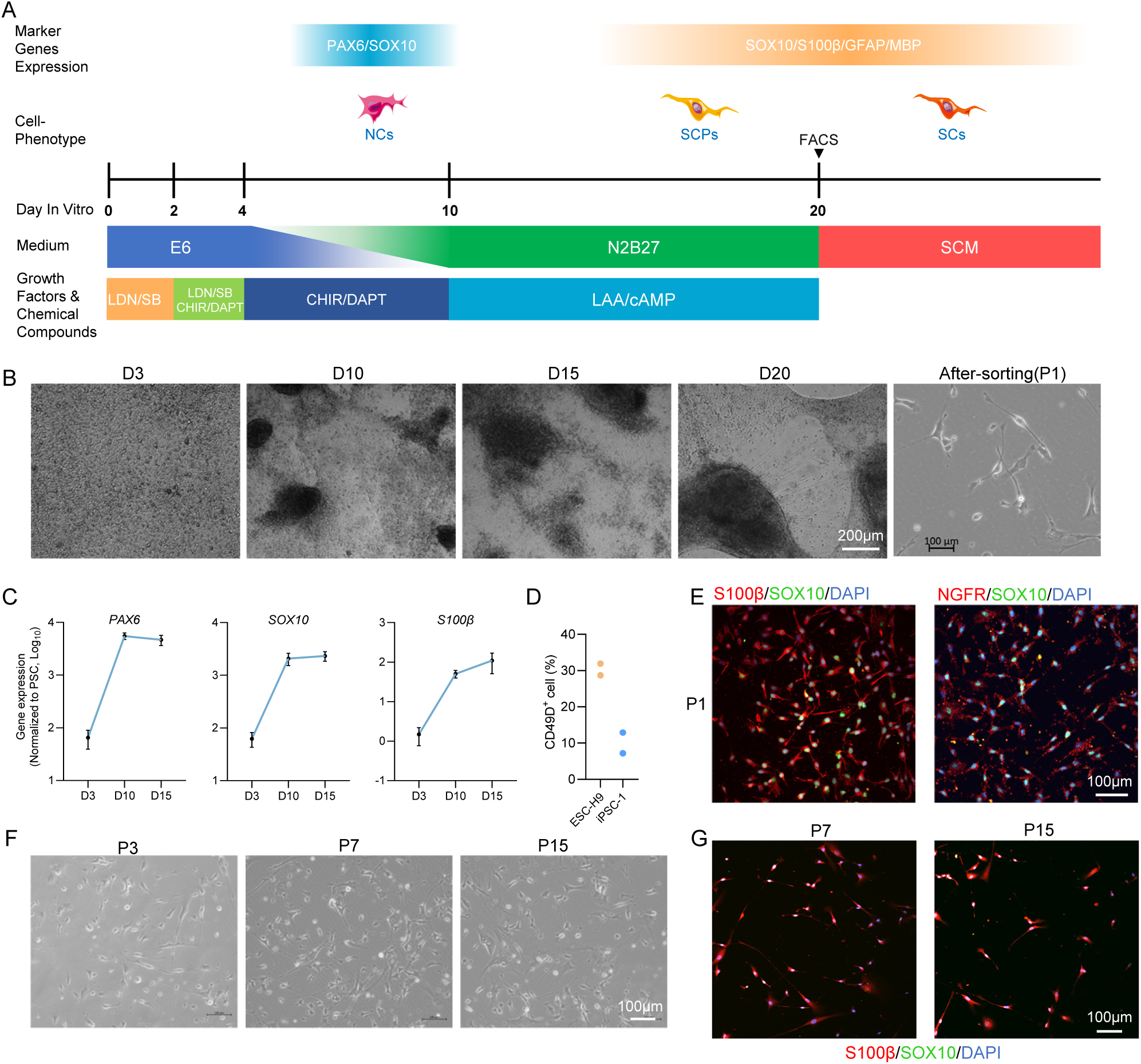
Efficient derivation, purification, and long-term expansion of Schwann cells from hPSCs. **A.** Schematic illustrating the differentiation conditions used to obtain schwann cells (SCs). NCs: Neural Crest cells; SCPs: Schwann Cell Progenitors. **B.** Bright-field images during the differentiation process. **C.** Gene expression analysis of SCs and progenitor cell markers of in vitro differentiation in 2 independent differentiations, data are normalized to that of PSC and shown as mean ± SD (n = 4 biological replicates per time point, N = 8 total; one-way ANOVA: F (2, 21) = 127; P < 0.001 for *PAX6*, F (2, 21) = 69.8; P < 0.001 for *SOX10*, F (2, 21) = 19.8; P < 0.001 for *S100β*). **D.** Efficiency of differentiation from 2 cell lines. **E.** Immunofluorescence staining showing the expression of SCs markers (SOX10, S100β, and NGFR). **F-G.** Bright-field and IF staining images of SCs during high-generation expansion.

### 2. Assembly of peri-nervoid and establishment of a transection injury model

Having established the three cellular building blocks, we next assembled them into a three-dimensional construct—the “Peri-nervoid” (PNO)—designed to recapitulate the cellular complexity of native peripheral nerves.

To construct Peri-nervoids (PNOs) recapitulating native peripheral nerve architecture, we assembled PNOs using a transparent silicone tube (inner diameter: 1.5 mm) filled with type I collagen gel (2 mg/mL) as the supporting matrix. SCs were uniformly resuspended in the gel at 3,000 cells/μL. A single mature SCO (D45, containing motor neurons) or 2-3 DRGOs (D45, containing sensory neurons, pooled to match the volume of one SCO) were placed at the center of the tube lumen, embedded within the SCs-laden collagen gel (Fig. 4A). Immediately after assembly (0 DPA), SCs were uniformly distributed within the gel with the organoid spheroid positioned at the center of the tube (Fig. 4B). By 14 DPA, abundant neurite bundles extended bidirectionally from the organoid spheroid along the longitudinal axis of the tube, traversing the entire gel and forming a macroscopically visible white cord-like structure (Fig. 4C). AAV-mediated labeling (SCO-GFP, DRGO-mCherry) exhibited motor (green) or sensory (red) axons forming mixed fascicles with interlaced spatial distribution (Fig. 4D). Immunofluorescence of PNO longitudinal sections at 14 DPA showed TUJ1⁺ axons extending along the tube axis with S100β⁺ SCs closely ensheathing the bundles and elongating in parallel, recapitulating the canonical axon-SCs aligned architecture (Fig. 4E–G). Axon outgrowth routinely exceeded 1 cm in length. Whole-mount imaging further showed highly organized fascicular axon bundles with SCs longitudinally apposed along their surfaces (Fig. 4H); Imaris colocalization analysis confirmed close spatial association between TUJ1⁺ axons and S100β⁺ SCs (white signals, Fig. 4h’).

**Fig. 4.**
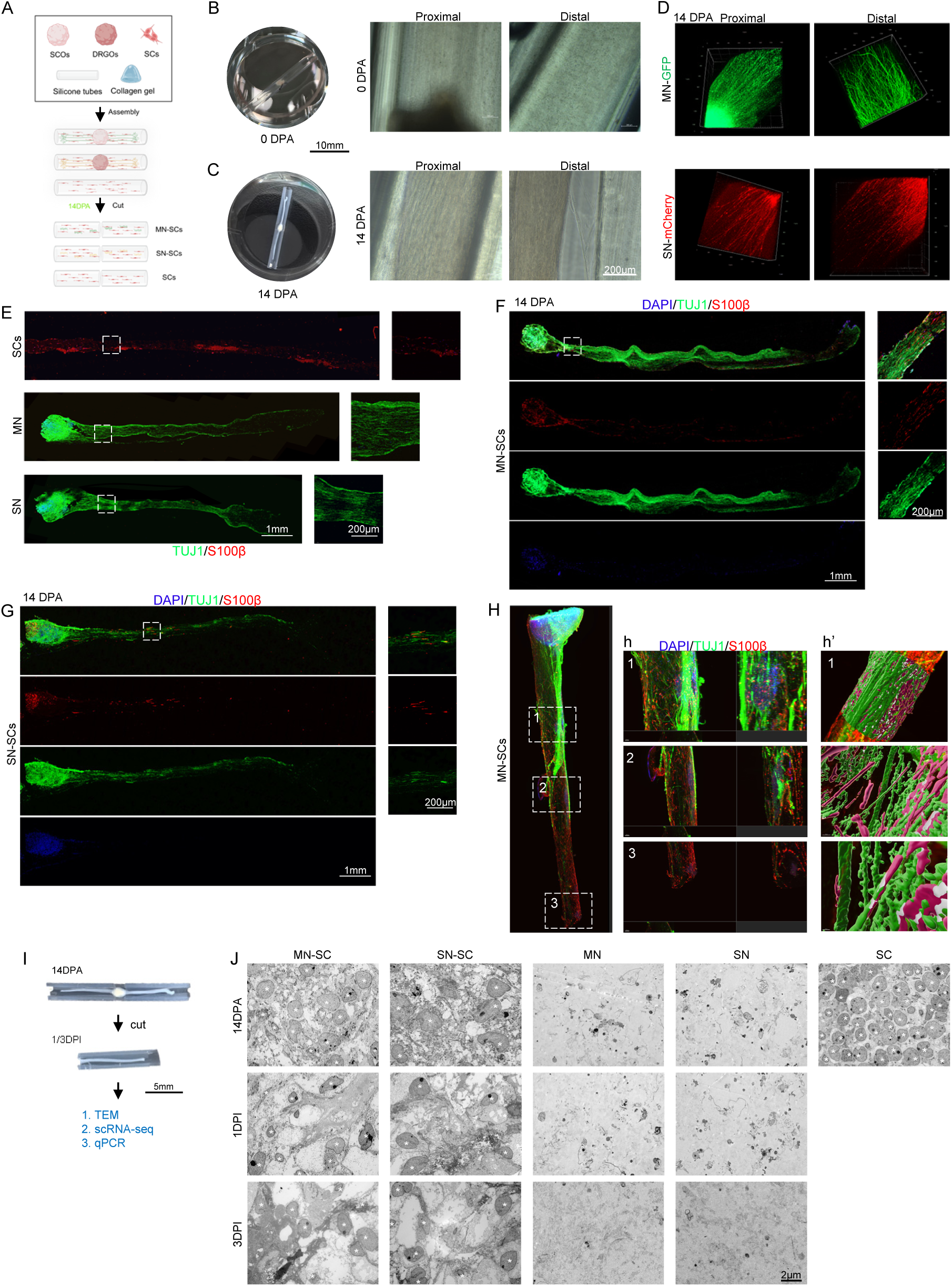
Three-dimensional assembly of Peri-nervoid recapitulating axon-SCs aligned architecture and establishment of a transection injury model that reproduces early axonal degeneration. **A.** Schematic of peri-nervoid (PNO) fabrication. SCOs (motor neurons), DRGOs (sensory neurons), and Schwann cells were assembled for 14 days (14 DPA), followed by removal of the spheroids to obtain the PNOs. Created in https://BioRender.com. **B-C.** Representative gross and bright-field images of PNOs at day 0 and day 14 of assembly. **D.** Three-dimensional view of axonal growth patterns of fluorescently labeled motor (green) and sensory (red) neurons. **E**. Immunofluorescence staining of SCs monocultures (S100β, red) and motor/sensory neurons monocultures (TUJ1, green) at day 14. The dashed box in the left panoramic view is shown at higher magnification on the right. **F-G.** Longitudinal cryosections of MN/SN-SCs-PNOs at 14 DPA immunostained for TUJ1 (green) and S100β (red). **H.** Whole-mount 3D light-sheet imaging of an intact PNO immunolabeled for TUJ1 (green) and S100β (red) after 3DISCO clearing. The white dashed boxes in H are shown at higher magnification (frontal and lateral views) (h). White signals represent colocalization of TUJ1 and S100β as calculated by Imaris colocalization analysis(h’).

To model the axonal injury that occurs during autologous nerve transplantation, we established a PNO transection injury model by severing the organoid spheroids from the extending axon bundles at 14 DPA (Fig. 4I). TEM analysis at 14 DPA (pre-injury) observed that SCs processes closely surrounded individual axons, without concentric lamellar myelin compaction (Fig. 4J). SCs nuclei (indicated by asterisks) displayed normal morphology with evenly distributed chromatin, and axons exhibited intact cytoskeletal ultrastructure. At 1 DPI, prominent axonal degeneration was evident, including increased axoplasmic density, microtubule/microfilament disintegration, loose and distorted myelin lamellae, and chromatin margination in SCs nuclei. By 3 DPI, further axonal vacuolization and phagocytic clearance of myelin debris by activated SCs were observed, with phagolysosomes containing myelin remnants within SCs cytoplasm. These results demonstrate that our PNO system successfully recapitulates native axon-SCs architecture with centimeter-scale outgrowth, and the transection model effectively reproduces early axonal degeneration and SCs responses following nerve injury, providing a robust in vitro platform for subsequent in vivo studies.

### 3. Single-cell transcriptomic profiling reveals context-dependent Schwann cell heterogeneity and injury-induced repair signatures that recapitulate Wallerian degeneration

Having established the PNO assembly and injury model, we next sought to characterize the transcriptional landscape of SCs within these constructs under different co-culture conditions and injury states, to understand how axonal signals and injury modulate SCs phenotypes. To this end, we performed scRNA-seq on five experimental groups: SCs cultured alone (sc), motor neuron-derived PNOs at 14 DPA (mn14dpa), sensory neuron-derived PNOs at 14 DPA (sn14dpa), motor neuron-derived PNOs at 3 days post-injury (mn3dpi), and sensory neuron-derived PNOs at 3 days post-injury (sn3dpi). After quality filtering, we obtained a total of 58,943 cells across all samples. UMAP dimensionality reduction and unsupervised clustering identified seven transcriptionally distinct SCs subclusters, which were annotated based on the expression of canonical SCs subtype markers (Fig. 5A-D, Supplementary Fig. 1, Table 1). These included repair-associated SCs (Repair-rSCs), myelinating SCs (mySCs), proliferating SCs (proSCs), lipid handling, and immature/precursor populations (Fig. 5C-D, Supplementary Fig. 1). Marker gene analysis confirmed that Repair-rSCs were characterized by the selective upregulation of AREG, SAA1, and BIRC3, whereas mySCs expressed high levels of PPARG, SGCD, and PTGS2 (Fig. 5C, G, Supplementary Fig. 1). proSCs were distinguished by the expression of proliferation-associated genes including PRPH and CCNA2 (Fig. 5C, Supplementary Fig. 1).

**Fig 5.**
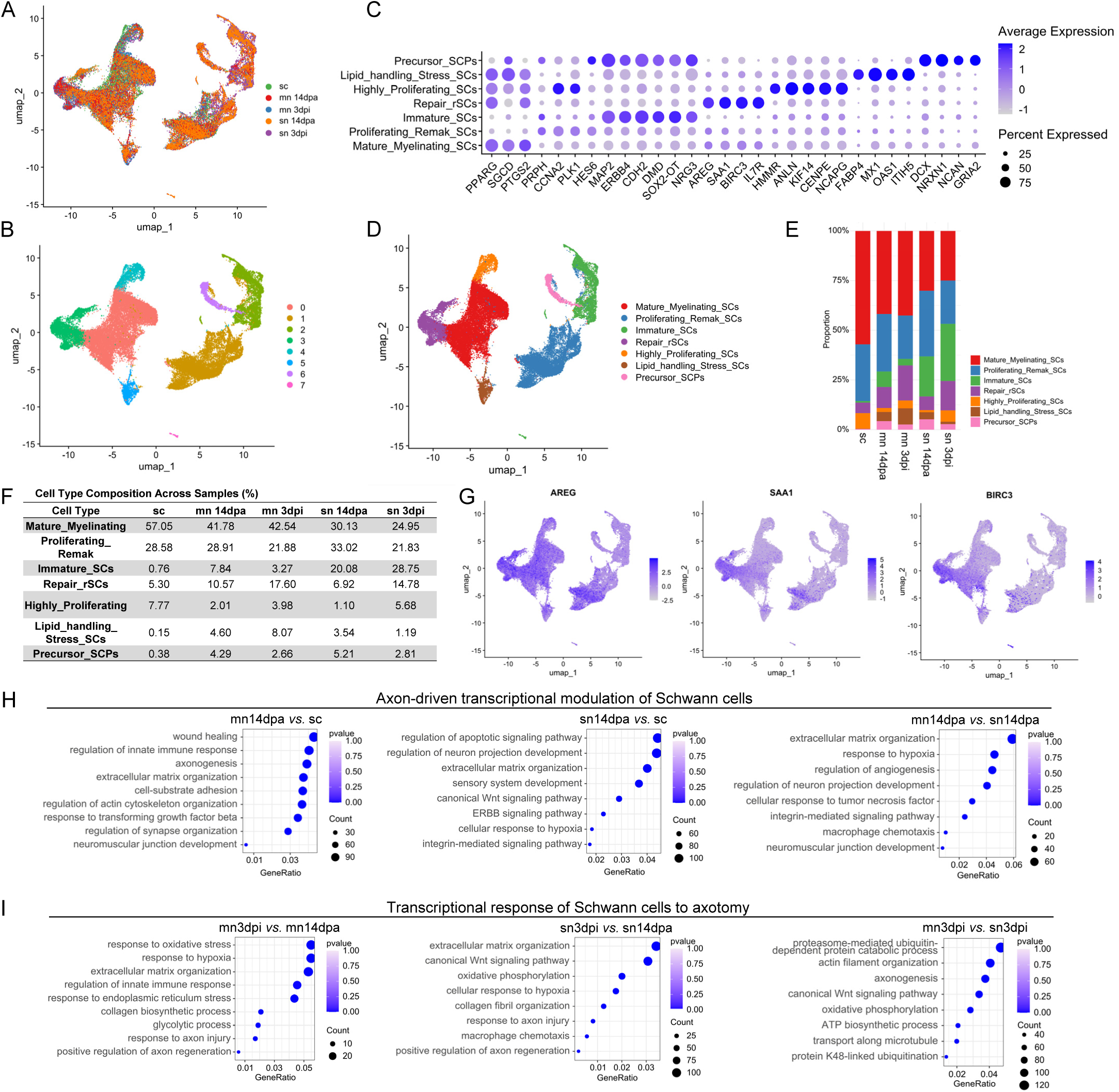
Singlecell RNAsequencing profiling reveals the heterogeneity and transcriptional dynamics of Schwann cells across different coculture conditions and injury states. **A.** UMAP projection of all single cells, colored by sample origin: sc (Schwann cells alone), mn14dpa (motor neuron PNOs at 14 DPA, preinjury), mn3dpi (motor neuron PNOs at 3 days postinjury), sn14dpa (sensory neuron PNOs at 14 DPA, preinjury), and sn3dpi (sensory neuron PNOs at 3 days postinjury). **B.** UMAP plot showing unsupervised clustering of cells based on transcriptomic similarity, with distinct clusters colorcoded. **C.** Dot plot depicting the expression of canonical Schwann cell subtype marker genes across identified clusters. Dot color indicates average expression level; dot size represents the percentage of cells expressing the gene within each cluster. **D.** UMAP projection annotated with the seven transcriptionally distinct Schwann cell subclusters. **E-F.** Stacked bar plot (E) and summary table (F) showing the proportional shift in SC subcluster composition across conditions. G. Feature plots illustrating the expression of Repair Schwann cell (Repair_rSCs) specific marker genes *AREG*, *SAA1*, and *BIRC3*, highlighting their selective enrichment in distinct subpopulations. **H-I.** Bubble plots of Gene Ontology (GO) biological process enrichment analysis specifically performed on differentially expressed genes (DEGs) identified within the Repair-rSCs subcluster across pairwise sample comparisons. The xaxis represents GeneRatio (the proportion of DEGs in a given GO term among all DEGs); bubble size reflects the number of enriched genes; bubble color indicates statistical significance (adjusted pvalue).

Strikingly, the proportional composition of these subclusters varied markedly across experimental conditions (Fig. 5E-F). In SC-alone cultures (sc), the majority of cells corresponded to myelinating/ proliferating populations, with minimal representation of repair or precursor subtypes. In contrast, co-culture with either motor (mn14dpa) or sensory (sn14dpa) neurons shifted the SC transcriptional landscape toward greater cellular diversity, while myelinating SCs (mySC) remained the dominant population, the proportions of precursor, immature, repair, and lipid-handling subclusters increased, accompanied by a marked decline in the proliferating SC population. The proportion of Repair-rSCs was dramatically increased following axonal transection (mn3dpi and sn3dpi), rising from less than 10% in pre-injury samples to over 15% at 3 DPI, accompanied by a corresponding decline in precursor subclusters. These injury-induced reprogramming—loss of myelinating phenotype, upregulation of repair and immature markers, and emergence of Repair-rSCs—recapitulate the core features of Schwann cell de-differentiation observed during Wallerian degeneration in *vivo*, where SCs rapidly switch from a myelinating to a repair-supportive state upon loss of axonal contact. This injury-induced shift was observed in both motor and sensory PNO contexts, indicating that the SC response to axonal injury is largely conserved across neuronal subtypes. However, subtle differences were noted between motor and sensory groups: the proportion of Repair-rSCs in mn3dpi exceeded that in sn3dpi, while mySC depletion was more pronounced in the sensory injury group (Fig. 5F). To validate the transcriptional reprogramming of SCs observed in our scRNA-seq analysis, we performed qPCR on PNOs at 14 DPA (co-culture with axons) and 3 DPI (post-transection), using SCs cultured alone as the control (Supplementary Fig. 2). Compared with SCs cultured alone, all examined genes were significantly upregulated in both co-culture (14 DPA) and injury (3 DPI) conditions, indicating that axonal contact alone broadly activates SC transcriptional programs across precursor, myelinating, trophic, and repair-associated pathways. Comparison between 14 DPA and 3 DPI revealed distinct dynamic regulation of SC phenotypes following axonal transection. At 3 DPI, in both MN and SN groups, the expression of precursor/undifferentiation markers (*SOX1*, *SOX2*), neurotrophic factors (*NGF*, *NT3*), and Remak/non-myelinating markers (*GFAP*) were upregulated compared to 14 DPA, reflecting a shift toward a repair-associated and less mature state. Conversely, myelination markers (*MPZ*, *P*MP22, *EGR2*, and *GPR126*) were downregulated at 3 DPI relative to 14 DPA. Together, these findings suggest that the PNO transection model faithfully reproduces the early phase of Wallerian degeneration in *vitro*.

To further characterize the functional properties of Repair-rSCs and their response to axonal injury, we performed Gene Ontology (GO) enrichment analysis on differentially expressed genes (DEGs) identified within the Repair-rSCs subcluster across pairwise comparisons: mn14dpa vs. sc, sn14dpa vs. sc, mn14dpa vs. sn14dpa, mn3dpi vs. mn14dpa, and sn3dpi vs. sn14dpa (Fig. 5H-I). In mn14dpa vs. sc and sn14dpa vs. sc comparisons, DEGs were significantly enriched for extracellular matrix organization, TGF-β signaling, and cell adhesion, indicating that axonal contact drives SCs toward a matrix-remodeling phenotype. The mn14dpa vs. sn14dpa comparison indicated enrichment of pathways related to neurotrophic signaling and ion transport, pointing to subtype-specific regulatory differences between motor and sensory SCs environments. More importantly, the injury-associated comparisons (mn3dpi vs. mn14dpa and sn3dpi vs. sn14dpa) identified robust enrichment of biological processes associated with inflammatory response, glial activation, and regulation of axon regeneration, including terms such as “response to axon injury,” “response to hypoxia,” and “positive regulation of axon regeneration” (Fig. 5I).

Collectively, these data demonstrate that SCs undergo profound context-dependent transcriptional reprogramming. Co-culture with axons drives SCs toward a more diverse transcriptional landscape in which myelinating phenotypes remain dominant while precursor, immature, repair, and lipid-handling subpopulations expand at the expense of proliferating SCs. Axonal transection further shifts the balance, triggering a rapid and robust increase in Repair-rSCs alongside a concomitant decline in proliferating and myelinating populations, paralleled the canonical SC response during Wallerian degeneration in *vivo*. The presence of sensory versus motor axons imposes distinct transcriptional signatures, suggesting that SC plasticity is finely tuned by the neuronal subtype with which they interact, and that axonal injury further drives this phenotypic transition from myelinating to repair-oriented states.

### 4. MN-PNO and SN-PNO grafts exhibit comparable reparative efficacy, each outperforming SCs-alone grafts in axonal outgrowth, and human SC retention at both 14 and 70 days post-transplantation

Having established the molecular and cellular properties of PNOs in vitro, we next evaluated their reparative efficacy in vivo. We established a 7-mm sciatic nerve defect model in immunodeficient NSG mice, with animals randomly assigned to six groups: (1) autograft, (2) collagen gel-filled hollow conduit, (3) SCs suspended in collagen gel alone, (4) MN-SCs (motor PNO), (5) SN-SCs (sensory PNO), and (6) MIX (MN-PNO + SN-PNO combined). For PNO-containing groups (MN-SCs, SN-SCs, and MIX), two PNO strands were combined into a single bundle within a silicone conduit before implantation. In the MIX group, one MN-PNO and one SN-PNO strand were paired together to provide both motor and sensory axonal components within the same graft (Fig. 6A–B). At 14 days post-transplantation (14 DPT), longitudinal sections immunostained for NF (axons) and S100β (SCs) demonstrated that the MN-SCs, SN-SCs, and MIX groups exhibited comparable axonal penetration, with NF⁺ fibers extending through the proximal, middle, and distal segments and maintaining structural integrity across the 7-mm defect (white dashed boxes delineating graft boundaries) (Fig. 6C). In contrast, the SCs-alone group showed markedly less axonal outgrowth; however, all cell-containing groups remained below the level of the autograft group (Fig. 6D). HNA/S100β co-staining confirmed survival of transplanted human SCs in all cell-containing groups (Fig. 6E). Quantitative analysis across proximal, middle, and distal segments further confirmed that the MN-SCs, SN-SCs, and MIX groups exhibited similar HNA⁺ cell proportions across all segments, with no significant proximal-to-distal decline in any of the three groups (p > 0.05), in contrast to the SCs-alone group which showed the lowest retention across all segments (p < 0.01) (Fig. 6F, Supplementary Fig. 3).

**Fig. 6.**
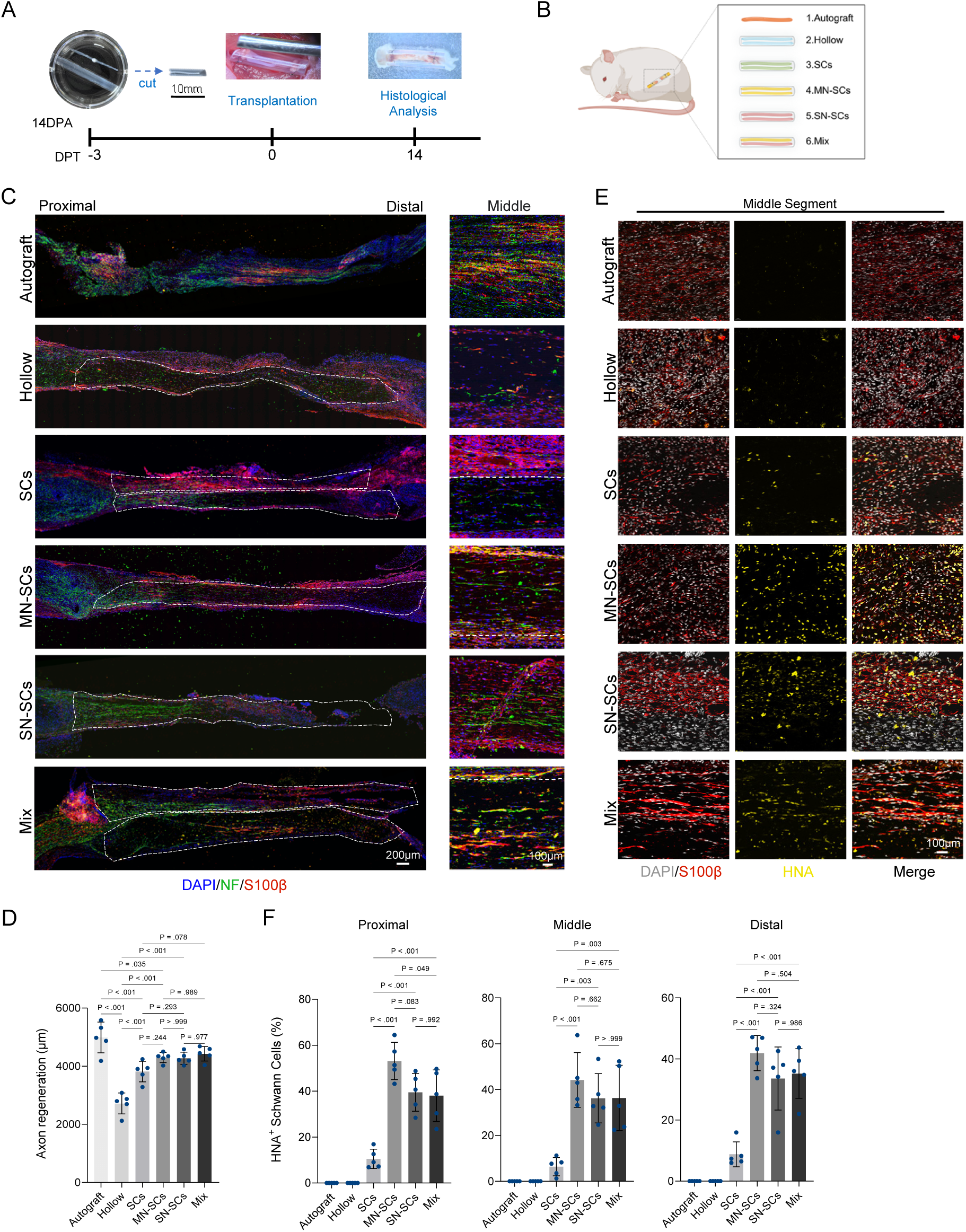
Early (14 DPT) in vivo evaluation demonstrates that PNO grafts promote axonal regeneration and human Schwann cell survival across the entire graft. **A.** Experimental timeline. **B.** Grouping: autograft, Hollow (silicone tube with collagen gel filled), SCs, MN-SCs, SN-SCs, and MIX; PNO grafts were assembled from two strands combined within a silicone tube. Created in https://BioRender.com. **C.** Longitudinal sections of grafts at 14DPT stained for NF (green) and S100β (red). The white dashed boxes delineate the identifiable boundaries of the PNO grafts within the host tissue. **D.** Quantification of axonal regeneration distance from (c) (n = 5 animals each group, one-way ANOVA followed by Tukey’s multiple-comparisons test: F (5, 24) = 26.4, P < 0.001). **E.** Mid-graft HNA (human nuclei, green) and S100β (red) co-staining. **F.** Quantification of HNA^+^ cell percentages across proximal, middle, and distal graft segments (n = 5 animals each group, one-way ANOVA followed by Tukey’s multiple-comparisons test: F (3, 16) = 23.0; P < 0.001 for Proximal, F (3, 16) = 11.6; P < 0.001 for Middle, F (3, 16) = 7.97; P = 0.003 for Distal). Data are shown as mean ± SD.

At 70 days post-transplantation (70 DPT), longitudinal sections confirmed long-term survival and widespread distribution of HNA⁺/S100β⁺ human SCs in all cell-containing groups, with the MN-SCs, SN-SCs, and MIX groups again exhibiting comparable proportions across all segments and no significant proximal-to-distal decline (p > 0.05). The SCs-alone group continued to show the lowest retention (Fig. 7B–C). Cross-sectional analysis across the MN-SCs, SN-SCs, and MIX groups showing similar proportions of human SCs among total MBP⁺ cells, significantly surpassing the SCs-alone group (p < 0.05), and displaying dense, homogeneous HNA⁺/MBP⁺ cell distribution (Supplementary Fig. 4. A–B).

**Fig. 7.**
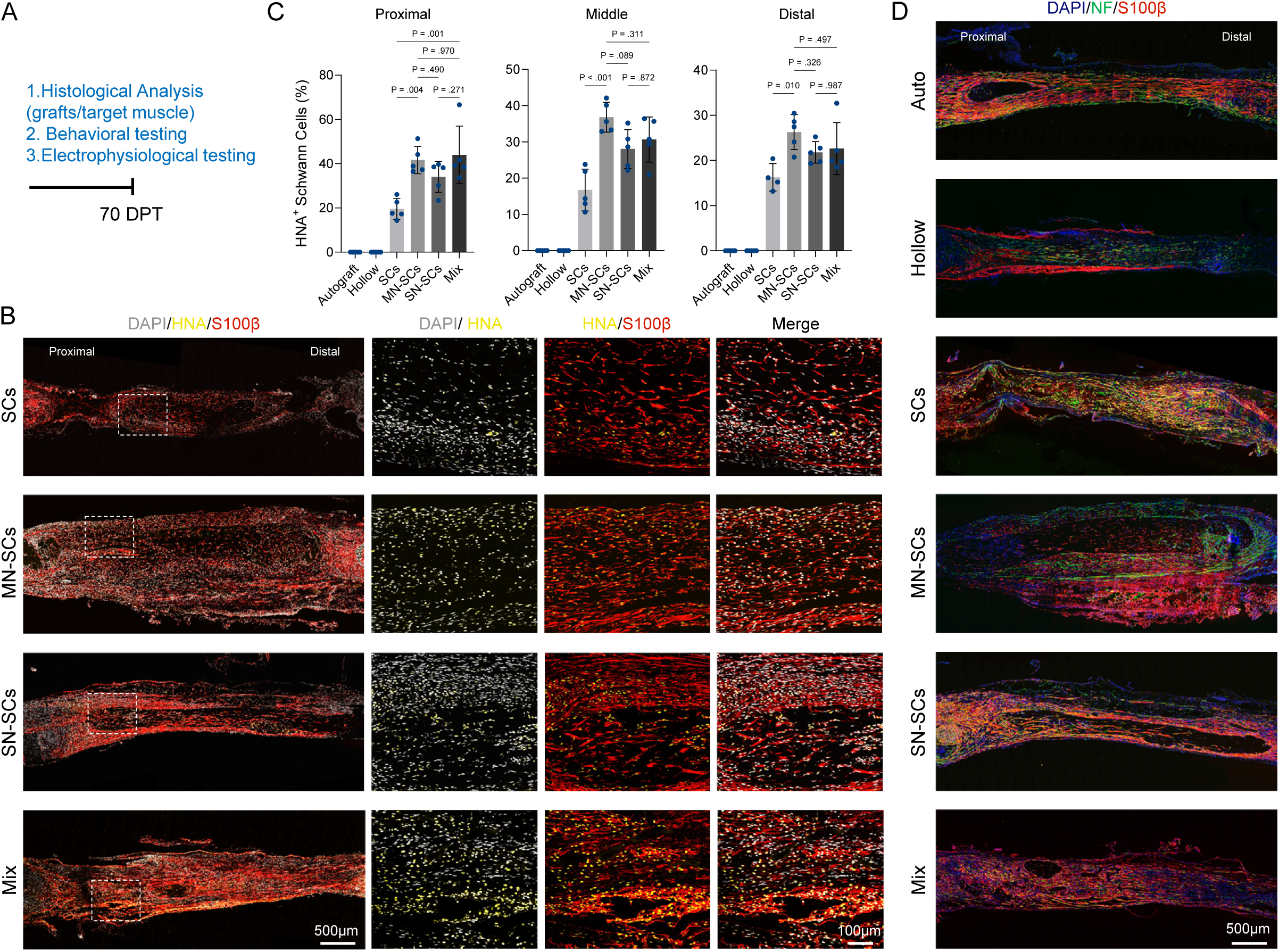
Long-term (70 DPT) in vivo evaluation confirms sustained axonal regeneration and persistent human Schwann cell engraftment in PNO grafts. **A.** Experimental timeline showing endpoint analyses (histology, behavior, electrophysiology). **B.** Longitudinal graft sections stained for HNA (green) and S100β (red); white dashed boxes indicate regions shown at higher magnification (right). **C.** Quantification of HNA^+^ cell percentages in proximal, middle, and distal segments (n = 5 animals each group, one-way ANOVA followed by Tukey’s multiple-comparisons test: F (3, 16) = 8.71; P = 0.001 for Proximal, F (3, 16) = 12.0; P < 0.001 for Middle, F (3, 15) = 4.63; P = 0.018 for Distal). **D.** Longitudinal graft sections stained for NF (green) and S100β (red). Data are shown as mean ± SD.

Longitudinal NF/S100β staining at 70 DPT showed that the MN-SCs, SN-SCs, and MIX groups all exhibited robust axonal regeneration, with NF⁺ fibers spanning the entire graft and S100β⁺ SCs closely ensheathing axon bundles in characteristic aligned architecture; the SCs-alone group showed markedly less axonal outgrowth (Fig. 7D). Cross-sectional analysis confirmed comparable axonal density (NF⁺ area percentage) among the MN-SCs, SN-SCs, and MIX groups, all approaching autograft levels (Supplementary Fig. 4. C–D).

These long-term histological findings demonstrate that grafts containing either MN-SCs, SN-SCs, or both in combination all promote superior axonal regeneration, remyelination, and human cell retention compared to SCs alone, with no significant differences among the three neuronal-containing configurations.

### 5. Histological, Ultrastructural, and Functional Outcomes at 70 days post-transplantation

We next assessed the quality of nerve regeneration and functional recovery at 70 days post-transplantation through comprehensive histological, ultrastructural, and functional evaluations. Toluidine blue staining of mid-graft cross-sections showed abundant, organized myelinated nerve fibers in all cell-containing groups (SCs, MN-SCs, SN-SCs, and MIX), comparable to the autograft, whereas the hollow conduit showed sparse, disorganized fibers (Fig. 8A). TEM ultrastructural analysis confirmed well-compacted myelin sheaths and normal axoplasmic ultrastructure in all cell-containing groups, resembling the autograft, while the hollow conduit exhibited thin, loosely packed myelin with axonal degeneration features (Fig. 8B). Quantitative analyses demonstrated that all cell-containing groups achieved comparable myelin sheath thickness (Fig. 8F) and optimal G-ratio values (∼0.65–0.70), all approaching autograft levels and indicating high-quality myelination, with no significant differences among the four groups (Fig. 8G).

**Fig. 8.**
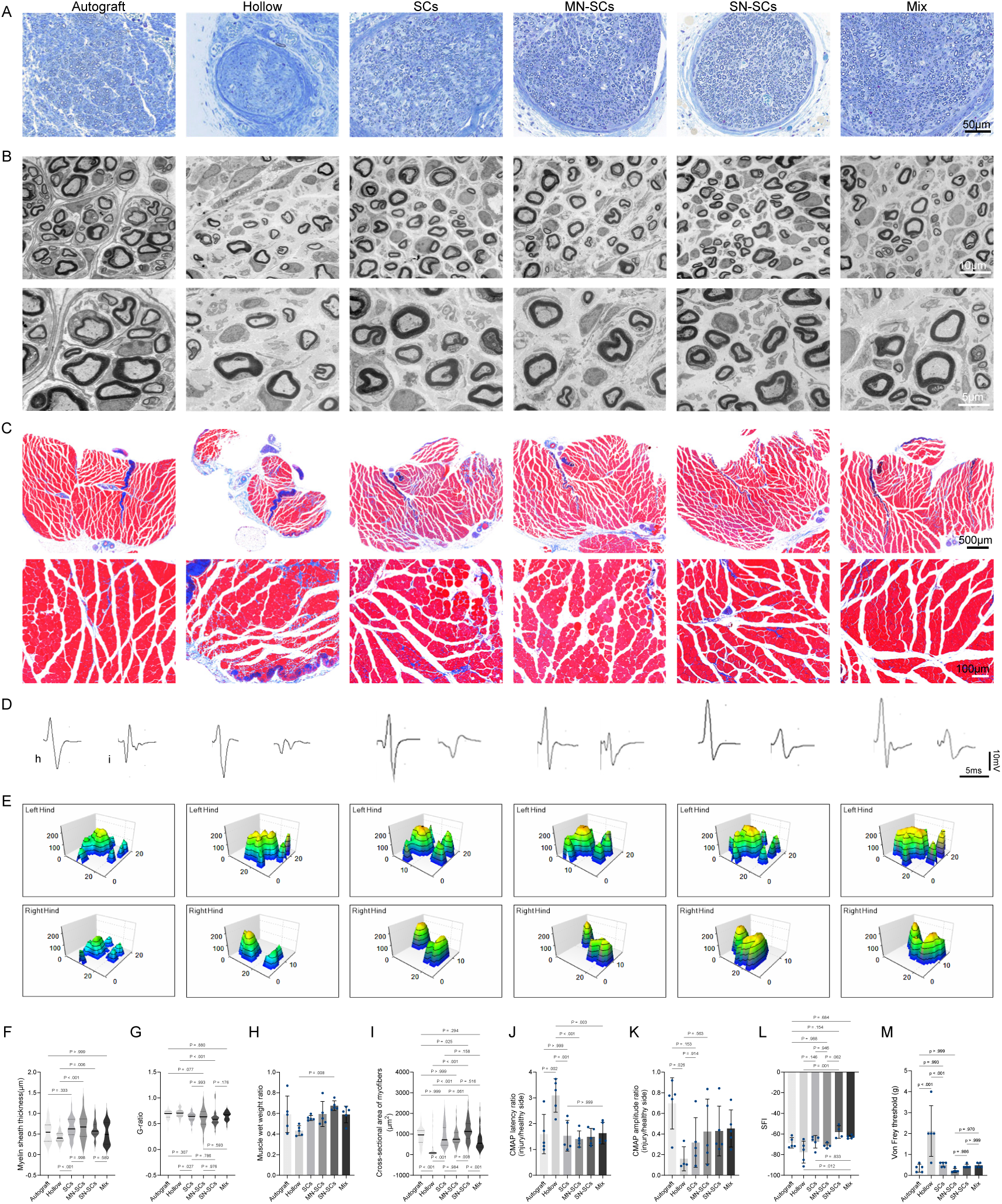
All cell-containing grafts achieve high-quality myelination and functional recovery comparable to autografts. **A.** Toluidine blue-stained cross-sections of grafts at mid-level. **B.** TEM images of the mid-graft; lower panels are magnified views of the boxed areas in upper panels. **C.** Masson’s trichrome staining of gastrocnemius muscles; lower panels show higher magnification. **D.** Representative EMG waveforms from healthy contralateral (h) and injured ipsilateral (i) sides. **E.** Catwalk 3D paw print images. **F-I.** Quantification of myelin sheath thickness (F), G-ratio (G), muscle wet weight recovery ratio (H), and mean muscle fiber area (I) (one-way ANOVA followed by Tukey’s multiple-comparisons test: F (5, 227) = 6.00; P < 0.001 for F, F (5, 230) = 5.79; P < 0.001 for G, F (5, 24) = 3.26; P = 0.022 for H, F (5, 287) = 13.3; P < 0.001 for I). **J-K.** Quantification of CMAP latency ratio (J) and amplitude ratio (K) (one-way ANOVA followed by Tukey’s multiple-comparisons test: F (5, 24) = 7.5; P < 0.001 for J, F (5, 23) = 2.56; P = 0.055 for K). **L.** SFI values from Catwalk analysis (F (5, 22) = 5.40; P = 0.002). **M.** Quantification of von Frey threshold (F (5, 24) = 9.73; P < 0.001). Data are shown as mean ± SD, n = 5 animals each group.

Masson’s trichrome staining of gastrocnemius muscles from the injured hindlimb exhibited well-preserved muscle architecture with organized myofibers and minimal collagenous deposits in all cell-containing groups and the autograft group, in contrast to extensive fibrotic infiltration and severe atrophy in the hollow conduit group (Fig. 8C). Quantitative analysis confirmed that all cell-containing groups achieved significantly higher wet weight recovery ratios (Fig. 8H) and larger mean muscle fiber cross-sectional areas (Fig. 8I) compared to the hollow group (p < 0.01), approaching autograft levels with no significant differences among the four groups, confirming effective attenuation of denervation-induced atrophy.

Electrophysiological evaluation at 70 DPT showed that CMAP amplitude recovery of all cell-containing groups approaching autograft values with no significant differences among the four groups (Fig. 8K). CMAP latency ratios showed no significant differences between any cell-containing group and the autograft group (p > 0.05), while the hollow conduit exhibit prolonged latency (p < 0.05) (Fig. 8J). Representative CMAP waveforms demonstrated robust action potential amplitudes in all cell-containing and autograft groups, contrasting with markedly attenuated responses in the hollow conduit group (Fig. 8D).

CatWalk gait analysis indicated that all cell-containing groups exhibited three-dimensional plantar pressure distribution patterns comparable to the autograft, while the hollow conduit showed severely impaired weight-bearing capacity (Fig. 8E). The sciatic functional index (SFI), a comprehensive measure of motor recovery, with values near 0 indicating normal function and near -100 indicating severe dysfunction, at 70 DPT was −68.58 ± 4.98 (autograft), −62.84 ±1.57 (MIX), −58.05 ± 6.10 (SN-SCs), −71.18 ± 2.48 (MN-SCs), −67.51 ± 6.09 (SCs), and −77.56 ± 10.61 (hollow). All cell-containing groups showed no significant difference from the autograft, and higher SFI values compared to the hollow group (p < 0.05) (Fig. 8L).

Sensory function was assessed by von Frey filament testing for mechanical allodynia and hot/cold plate tests for thermal sensation. In the von Frey test (Fig. 8M), the MN-SCs group exhibited significantly lower withdrawal thresholds compared to all other groups, including the autograft group. No significant differences were observed among the SCs, SN-SCs, MIX, and autograft groups. In contrast, hot and cold plate tests exhibited no significant differences in thermal withdrawal latencies between any of the cell-containing groups and the autograft group, suggesting comparable thermal sensory recovery across all groups (Supplementary Fig. 5).

Collectively, these combined ultrastructural, electrophysiological, and functional assessments demonstrate that all cell-containing grafts promote superior nerve regeneration, high-quality myelination, and target muscle reinnervation, achieving outcomes comparable to autologous nerve transplantation. These findings validate the multidimensional biomimetic “Peri-nervoid” assembly strategy as a promising therapeutic approach for peripheral nerve repair.

### 6. Biosafety assessment confirms the absence of abnormal proliferation, systemic toxicity, and organ damage following Peri-nervoid transplantation

To comprehensively evaluate the in vivo biosafety of the Peri-nervoid grafts, we performed systematic histological, hematological, and serum biochemical analyses at 70 DPT.

First, we assessed the potential risk of abnormal proliferation of transplanted human Schwann cells. Immunofluorescence co-staining for HNA (human nuclei) and Ki67 (proliferation marker), followed by quantitative analysis, was performed on samples collected at 14 DPA (baseline, pre-injury), 3 DPI (post-transection), and 70 DPT (Supplementary Fig. 6). The percentage of HNA⁺/KI67⁺ double-positive cells remained consistently low (∼8.67%) across all time points and experimental groups, with no significant differences detected among groups (p > 0.05), indicating that transplanted human Schwann cells did not undergo abnormal proliferation or neoplastic transformation during in vitro assembly, following injury stimulation, or after long-term in vivo engraftment (Supplementary Fig. 6).

Next, histopathological evaluation of grafts and major organs was performed via H&E staining. At 70 DPT, H&E staining of mid-graft sections from all groups showed normal tissue architecture with no evidence of tumor formation, hypercellularity, or atypical cell clusters (Supplementary Fig. 7A). H&E staining of major organs, including heart, liver, spleen, lung, and kidney, also showed no signs of inflammatory infiltration, necrosis, or tumorigenesis across any of the experimental groups (Supplementary Fig. 7B).

Furthermore, systemic toxicity was evaluated through complete blood count and serum biochemistry analysis. No significant differences were observed in any hematological or serum biochemical parameters among the experimental groups (p > 0.05), further excluding potential impacts of the grafts on hematopoietic and hepatorenal functions (Supplementary Fig. 7C, D).

Collectively, the histopathological, hematological, and serum biochemical evaluations consistently demonstrated that Peri-nervoid grafts did not induce abnormal cell proliferation, systemic toxicity, or major organ damage, confirming the favorable biosafety profile of the grafts in recipient animals.

## Discussion

Previous studies have shown that hiPSC-derived SCs resemble primary adult SCs at the transcriptomic, proteomic and functional levels and can support peripheral nerve regeneration^15,23^. Our study extends these works from cell delivery to tissue organization. We combined hPSC-derived SCs with motor, sensory or mixed organoid-derived axons within collagen-filled silicone conduits to generate preformed nerve-like grafts. Following transplantation, human SCs survived throughout the graft for up to 10 weeks and contributed to the myelination of regenerating host axons. The SC-only and axon-containing grafts all supported axonal regeneration, myelination and motor recovery, with outcomes comparable to those of autografts at the assessed time points. These findings show that hPSC-derived SCs can retain their regenerative function within a structured three-dimensional graft rather than being delivered solely as a cell suspension.

Single-cell RNA sequencing further showed that the transcriptional state of SCs was shaped by both axonal context and injury. Co-culture with intact axons promoted a myelination-associated programme, whereas axonal transection induced rapid downregulation of myelination genes and upregulation of repair-associated and neurotrophic genes. This transition was consistent with the injury-induced conversion of differentiated SCs into repair SCs during Wallerian degeneration. SCs co-cultured with motor axons displayed a more pronounced injury-responsive signature than those co-cultured with sensory axons, suggesting that axonal identity may influence the SC response to injury. However, this transcriptional difference was not accompanied by detectable differences in functional recovery among the axon-containing grafts, and its biological significance therefore requires further investigation.

The biosafety of Peri-nervoid grafts was systematically evaluated through multiple approaches. HNA/Ki67 co-staining showed consistently low levels of proliferating human cells across all time points and groups, with no significant differences among experimental conditions. H&E staining of grafts and major organs revealed no evidence of inflammation, necrosis, or tumor formation, while all hematological and biochemical parameters remained within normal ranges with no significant differences among groups. These observations are particularly relevant given the tumorigenic risk associated with residual undifferentiated pluripotent stem cells^24^; FACS purification of CD49d^+^ SCs and removal of organoid spheroids before transplantation may have reduced this risk. Although longer follow-up and evaluation in immunocompetent and large-animal models remain necessary, these findings provide initial evidence for the biosafety of Peri-nervoid grafts under the conditions examined and support their further preclinical development.

The use of NSG mice enabled sustained engraftment of human cells by markedly reducing xenogeneic immune rejection, but it also precluded evaluation of graft performance in an intact immune system. Wallerian degeneration involves coordinated responses among SCs, immune cells and other non-neuronal populations^6^. Previous studies have demonstrated that immune cells, particularly macrophages and T cells, play critical roles in peripheral nerve regeneration^2,25^. Our findings in NSG mice may therefore not fully predict either the regenerative efficacy or the immunogenicity of Peri-nervoid grafts in immunocompetent recipients. For clinical translation, patient-specific hiPSC-derived SCs may reduce alloimmune responses but would require individualized manufacturing, long preparation times and substantial costs. Allogeneic hiPSC-derived SCs could offer an off-the-shelf alternative, although their immunogenicity and the need for immunosuppression remain to be established. Gene-edited hypoimmunogenic hiPSC lines may provide another strategy to reduce immune recognition^26^, but their long-term safety, phenotypic stability and regenerative efficacy require further investigation.

Several additional limitations should be acknowledged. First, the hPSC-SCs used in this study were Several limitations should be acknowledged. First, the hPSC-SCs used in this study were FACS-purified CD49d^+^ cells, representing a relatively differentiated stage. Schwann cell precursors (SCPs) could be generated more rapidly, expanded more efficiently and may promote axonal ingrowth more effectively than differentiated SCs^27^. Direct comparisons across developmental stages are therefore needed to determine the optimal cell population for graft construction. Second, the 10-week period precluded assessment of long-term cell persistence, myelin stability and the durability of functional recovery. Third, while we demonstrated functional motor recovery through SFI and electrophysiology, the quality of reinnervation, including axon targeting accuracy and synaptic organization, requires further investigation through retrograde tracing and ultrastructural analysis. Scaling the Peri-nervoid to clinically relevant nerve defects presents additional challenges, particularly in vascularization and manufacturing precision. Vascularization remains a core bottleneck limiting the survival of large-segment grafts; future strategies could incorporate pre-vascularization approaches (such as co-assembling hPSC-derived endothelial cells) or controlled-release systems for pro-angiogenic factors to accelerate directional host vessel ingrowth. In addition, the current self-assembly method provides limited control over SC distribution and axonal orientation. Three-dimensional bioprinting and microfluidic guidance may improve spatial organization, batch-to-batch consistency and functional reproducibility. Addressing vascularization, internal architecture and manufacturing consistency will be necessary before the performance of Peri-nervoid grafts can be evaluated in larger defects and more clinically relevant models.

In conclusion, we developed the Peri-nervoid, a preformed nerve-like graft that integrates hiPSC-derived SCs with motor and sensory axonal components. Within this construct, SCs responded dynamically to axonal context and injury, survived after transplantation, and contributed to axonal regeneration and remyelination in a 7-mm sciatic nerve defect, with no evidence of abnormal proliferation or systemic toxicity during the 10-week period. By combining structural guidance with living, injury-responsive cellular components, the Peri-nervoid moves engineered nerve grafts beyond passive structural mimicry and provides a foundation for their further development in peripheral nerve repair.

## Methods

### Culture of hESC and hiPSC

Human embryonic stem cells (hESCs) / human induced pluripotent stem cells (hiPSCs) were used in line with the local ethical regulations. The stem cell line exhibited a normal karyotype as determined by genomic stability testing. Human ESC-H9 (DC-02) and iPSC-1 (RC01001-A, male, 36 years old, blood-derived) were cultured in a 6-well plate coated with hESC-Qualified Matrigel (Corning, #354277), without feeders, at 37°C, 5% CO_2_ environment using Essential 8 (E8, Gibco, #A1517001). The medium was changed daily and cells were passaged when they reached 90% confluency using the 0.5 mM EDTA. Mycoplasma testing was performed monthly on the culture plate. To minimize potential variations due to prolonged culturing, the hESC/hiPSC used throughout this study were all below passage 45. Cell lines used in each experiment are shown in Table 2.

### Generation of SCOs

The SCOs were generated from hESC/hiPSC as previously described^19,22^ with some modifications. Briefly, on day -1, confluent hESC/hiPSC were detached using Gentle Cell Dissociation Reagent (Gibco, #A11105-01) and further mechanically dissociated into single cells. Cells were resuspended and counted in E8 Medium containing 10 μM Y-27632 (MCE, #HY-10071) at a density of 9,000 cells per 100 μl. Next, 100μl cell suspension was added to each well of a 96-well ultra-low-attachment U-bottom plate (Corning, #7007). After 24 hours, on day 0, when embryoid bodies (EBs) had formed, 75 μl of E8 Medium was removed, and 100μl of Essential 6 (E6, Gibco, #A1516401) supplemented with 100 nM LDN-193189 (Sigma, #SML0559) and 10 μM SB-431542 (Sigma, #S4317), was added to each well. This medium was replaced daily within the first 6 days. From day 4 to day 20, the WNT activator CHIR-99021(3 μM, Sigma, #SML1046) was utilized for caudalization.

On day 6, spheroids were transferred to 24-well ultra-low-attachment plate (Corning, #3473) with B27 medium containing Neurobasal-A (Life Technologies, #10888022), β-mercaptoethanol (0.1 mM, Gibco, #21985023), 1×B-27 supplement without vitamin A (Life Technologies, #12587010), 1× GlutaMax (Gibco, #35050038), and supplemented with RA (0.1 μM, Sigma, #R2625), EGF (20 ng/ml, R&D Systems, #236-EG), FGF-2 (10 ng/ml, R&D Systems, #233-FB-025), and the SHH modulator smoothened agonist (SAG, 1 μM, Millipore, #566660) until day 20, with medium changes every 2 days.

From day 20 onward, organoids are transferred to the N2B27 medium, B27 medium supplemented with 1× N-2 supplement (Life Technologies, #17502048), BDNF (20 ng/ml, Peprotech, #450-02), IGF-1 (10 ng/ml, Peprotech, #100-11), L-Ascorbic Acid (AA, 200 nM, Wako, #321-44823) and cAMP (50 nM, Sigma, #D0627). The Notch pathway inhibitor DAPT (2.5 μM, STEMCELL technologies, #72082) was added for neuralization from day 20 to day 26. Mitomycin-C treatment (1µg/ml, MCE, HY-13316) was used once at day 26 for 2 h to reduce the non-neuronal population. Medium was changed every 3-4 days.

### Generation of DRGOs

The DRGOs were generated as previously described^28,29^ with some modifications. EB formation was in line with SCOs. On day 0, 75 μl of E8 Medium was removed, and 100μl of E6 medium supplemented with LDN-193189(100 nM) and SB-431542(10 μM), was added to each well. This medium was replaced daily within the first 3 days, phasing in N2B27 media beginning at day 4, and gradually increasing the amount to 100% by day 11.

From day 3 to day 7, combination of 5 inhibitors (5i), include LDN-193189(100 nM), SB-431542 (10 μM), CHIR-99021(3 μM), DAPT (2.5 mM) and SU5402(10 µM, Sigma, #SML0443) was utilized for further promote neural crest phenotypes. On day 7, the use of inhibitors LDN193189 and SB-431542 ceased, but CHIR-99021, DAPT, and SU-5402 continued to be added (3i).

From day 11 onward, the medium was changed to N2B27 with human-β-NGF (10 ng/ml, Peprotech, #450-01), BDNF (10 ng/ml, Peprotech, #450-02), NT-3 (10 ng/ml, Peprotech, #450-03), and GDNF (10 ng/ml, Peprotech, #450-10) for every 2-3 days. Mitomycin-C treatment (1 µg/ml) was used once at day 14 for 2h.

### Generation of Schwann cells (SCs)

The SCs were generated as previously described^30^ with some modifications. Briefly, on day-2, confluent hESC/hiPSC were resuspended and counted in E8 Medium containing 10 μM Y-27632 at a density of 30,000 cells per 500 μl. Next, 500 μl cell suspension were added to each well of a 24-well plate coated with Matrigel (354277). After 48 hours, on day 0, 500 μl of E6 Medium supplemented with LDN-193189(500 nM) and SB-431542(10 μM), was added to each well. On day 2, change medium containing SB-431542(10 μM), LDN-193189(500 nM), CHIR-99021(3 μM) and DAPT (10 μM), On day 4, the medium was changed containing CHIR-99021(3 μM) and DAPT (10 μM), phasing in N2B27 beginning at day 4, and gradually increasing the amount to 100% by day 10 with medium changed daily.

From day 10 onward, the medium was changed every other day with L-Ascorbic Acid (200 μM) and cAMP (200 μM) until day 20, when the cells are ready for purification.

### Schwann cell purification and expansion

Briefly, SCs were purified with Fluorescence Activated Cell Sorting (FACS) at day 20 of differentiation using a PE-conjugated antibody to the alpha4 integrin, CD49d (R&D Systems, FAB1354P). After FACS, SCs were replated onto Matrigel (354277) coated plates with Schwann Cell Medium (SCM, ScienCell, 1701), passaged every 3-4 days and maintained in culture for up to passage 15.

### Calcium imaging

The calcium flow of the organoids was monitored by incubating them with a calcium chemical indicator Fluo-4 AM (3 μM, Life Technologies, #C34852) for 30 min at 37°C. Subsequently, the organoids were washed three times with DPBS. Calcium images were captured at 1 frame per second for 100 seconds at 37°C using a high-content analysis system (PerkinElmer, Operetta CLS). Changes in fluorescence intensity were analyzed using ImageJ. The ΔF/F trajectory for each region of interest (ROI) was calculated by dividing the fluorescence over time by the baseline fluorescence. Signal decay was controlled by subtracting the mean baseline fluorescence. Signal attenuation was calculated by subtracting the average fluorescence of the background. ΔF/F = (FROI-Fb)/Fb, where F_ROI_ and Fb represent the average fluorescence value of ROI and whole frame, respectively.

### Multi-electrode array (MEA) recording

Organoids at D40 were plated on 6-well MEA plates allowed to achieve electrical stability. Subsequently, recordings were conducted in a Maestro MEA system using the AxIS Software Spontaneous Neural Configuration (Axion Biosystems). Using Axion Biosystems’ Neural Metrics Tool, active electrodes were set to more than 5 spikes/min. Bursts electrode were defined as inter-spike interval (ISI) threshold requiring minimally 5 spikes with a maximum ISI of 100 ms.

### Peri-nervoid (PNO) assembly

Silicone tubes (inner diameter: 1.5 mm, outer diameter: 2.0 mm; length: 15 mm) were sterilized prior to use. Type I collagen (5 mg/mL, Corning, #354236) was diluted with sterile 0.02 N acetic acid and 10× PBS to a final concentration of 2 mg/mL, and the pH was adjusted to 7.2–7.4 using 1 N NaOH. The collagen solution was maintained on ice to prevent premature polymerization.

Schwann cells (passage 3–5) were dissociated using Gentle Cell Dissociation Reagent, and resuspended in ice-cold 2 mg/mL type I collagen solution at a density of 3,000 cells/μL (3 × 10⁶ cells/mL). The cell–collagen suspension was gently mixed and maintained on ice.

Mature spinal cord organoids (SCOs, day 45, ∼400–600 μm in diameter) and dorsal root ganglion organoids (DRGOs, day 45, ∼200–300 μm in diameter) were collected using a wide-bore pipette tip. A single SCO or 2–3 DRGOs (pooled to match the total volume of one SCO) were selected per construct. Approximately 30 μL of the cell–collagen suspension was injected into the silicone tube lumen using a 26-gauge needle attached to a Hamilton syringe, avoiding air bubbles. The organoid spheroid(s) were gently placed at the center of the tube lumen using fine forceps under stereoscopic magnification. Assembled constructs were incubated at 37°C in a humidified 5% CO₂ incubator for 30–40 min to allow collagen polymerization. Following gelation, tubes were transferred to 6-well plates containing PNO maturation medium, N2B27 medium: SCM = 1:1, supplemented with BDNF, GDNF, and NT-3 (each at 10ng/ml). PNOs were maintained at 37°C in 5% CO₂ up to 14 days, with medium changes every 2–3 days.

For AAV-mediated labeling of axons, SCOs and DRGOs were infected prior to assembly with rAAV-hSyn-EGFP-WPRE-hGH-pA (for SCOs) or rAAV-hSyn-mCherry-WPRE-hGH-pA (for DRGOs) (Brainvta) at day 30 of differentiation, as previously described^22^. Briefly, organoid spheroids were incubated with viral solution (1.0 × 10¹¹ vg/mL) in 200 μL N2B27 medium overnight, followed by addition of fresh medium and further incubation. Fluorescence was observed 12 days post-infection.

### PNO transection injury model

To model axonal injury, PNOs cultured for 14 days (14 DPA) were subjected to spheroid removal. Under a stereomicroscope, the organoid spheroids were carefully severed from the extending axon bundles using a sterile 30-gauge needle and fine forceps. The severed spheroids were removed, leaving the axon bundles within the collagen gel. Injured PNOs were maintained in maturation medium and collected at 1 and 3 days post-injury (1 DPI and 3 DPI) for transmission electron microscopy, single-cell RNA sequencing, qPCR analysis and transplantation.

### Single-cell suspension preparation and scRNA-seq library construction

Collected PNOs were dissociated into single-cell suspensions under strict quality control parameters. Cell viability was assessed and required to be >85%, and the cell concentration was adjusted to an optimal range of 500-1,500 cells/μL. Single-cell transcriptomic libraries were constructed using the Chromium GEM-X Single Cell 3′ V4 Reagent Kits (10x Genomics, Pleasanton, CA, USA) according to the manufacturer’s protocol. Briefly, single cells were encapsulated into Gel Beads-In-Emulsion (GEMs) with barcoded gel beads and reverse transcription reagents using a “double-cross” microfluidic chip. Within the GEMs, cells were lysed, and mRNA was reverse-transcribed into barcoded cDNA. Following GEM breaking and cDNA amplification, libraries were fragmented, end-repaired, A-tailed, and ligated with sequencing adapters. The final library quality and fragment size distribution were validated before sequencing. The libraries were sequenced on an Illumina NovaSeq platform using a 150-bp paired-end (PE150) strategy, targeting a sequencing depth of approximately 120 Gb per sample to ensure robust transcriptome coverage.

### Primary data processing and quality control

Raw sequencing data (BCL files) were demultiplexed and processed using the Cell Ranger pipeline (10x Genomics). Reads were aligned to the human reference genome (GRCh38, 2024-A annotation) using the STAR algorithm. Unique Molecular Identifiers (UMIs) and cell barcodes were extracted, corrected for sequencing errors, and deduplicated to generate raw gene-barcode count matrices. Downstream computational analyses were performed in the R environment. To ensure high data integrity, rigorous quality control (QC) was applied to the filtered feature-barcode matrices. Cells were excluded if they fell outside the acceptable thresholds for total UMI counts (*nCount_RNA*), detected gene numbers (*nFeature_RNA*), or exhibited excessive mitochondrial gene expression (*percent.mt*), which is indicative of apoptotic cells or compromised membranes. Furthermore, to eliminate multiplet artifacts (doublets or multiplets) that could confound downstream clustering, we computationally identified and removed doublets using the scDblFinder R package, which simulates artificial doublets to train a robust classification model.

### Data integration, dimensionality reduction, and clustering

To construct a comprehensive transcriptomic landscape of iPSC-derived Schwann cells across five distinct sample conditions (designated as sc, sn 14dpa, sn 3dpi, mn 14dpa, and mn 3dpi), we employed the Seurat (v5) R package. Following standard log-normalization (NormalizeData) and the identification of the top 2,000 highly variable genes (FindVariableFeatures using the ’vst’ method), we mitigated batch effects across the five samples using Seurat’s Canonical Correlation Analysis (CCA)-based integration workflow. Integration anchors were identified (FindIntegrationAnchors) and used to generate a corrected, integrated expression matrix (IntegrateData) utilizing the first 20 principal components (PCs).

Dimensionality reduction was performed via Principal Component Analysis (PCA) on the integrated data. The top 20 PCs, determined by the JackStraw procedure and Elbow plot, were selected to construct a k-nearest neighbor (KNN) graph (FindNeighbors). Unsupervised graph-based clustering (FindClusters) was performed at varying resolutions to capture both broad lineage identities (resolution = 0.1) and fine-grained cellular heterogeneity (resolution = 0.5). The resulting clusters were visualized in two-dimensional space using Uniform Manifold Approximation and Projection (UMAP).

### Cell type annotation and Schwann cell subtyping

Cluster identities were annotated based on the expression of canonical Schwann cell lineage markers (e.g., *SOX10*, *S100B*, *MPZ*, *PLP1*, *ERBB3*, *SNAI2*) and validated using FeaturePlot and DotPlot visualizations. To dissect the functional heterogeneity within the Schwann cell lineage, sub-clustering was performed, revealing distinct developmental and reactive states. These subpopulations were annotated as: Mature Myelinating Schwann Cells (marked by *PPARG*, *SGCD*), Proliferating Remak Schwann Cells (*PRPH*, *CCNA2*), Immature Schwann Cells (*MAP2*, *ERBB4*), Repair Schwann Cells (rSCs; *AREG*, *SAA1*, *BIRC3*), Highly Proliferating SCs (*HMMR*, *ANLN*), Lipid-handling Stress SCs (*FABP4*, *MX1*), and Precursor Schwann Cell Precursors (SCPs; *DCX*, *NRXN1*). The proportional distribution of these subtypes across different sample conditions was quantified and visualized using stacked bar plots.

### Differential expression and functional enrichment analysis

To investigate the transcriptional reprogramming of Repair Schwann Cells (rSCs) across different microenvironments or injury models, differential expression analysis was conducted between predefined comparative groups (e.g., motor nerve vs. sensory nerve, injury vs. naive states) using the Wilcoxon rank-sum test implemented in the FindMarkers function. Genes were considered significantly differentially expressed (DEGs) if they met the following criteria: absolute log2 fold-change (|log2FC|) > 0.25, detected in a minimum of 10% of cells in either population (*min.pct* > 0.1), and an adjusted *P*-value < 0.01.

Prior to functional annotation, a stringent filtering step was applied to the DEG lists to exclude non-coding RNAs, pseudogenes, and uncharacterized transcripts (e.g., filtering out genes matching patterns such as *ENSG*, *LINC*, *MIR*, *-AS*, *-DT*, and *-PS*). Gene Ontology (GO) enrichment analysis for Biological Processes (BP) was performed using the clusterProfiler R package with the org.Hs.eg.db annotation database. Gene symbols were mapped to Entrez IDs, and *P*-values were adjusted for multiple testing using the Benjamini-Hochberg (BH) procedure. Enriched pathways with an adjusted *P* < 0.05 were considered significant and visualized using customized bubble plots.

### Statistical analysis and software availability

All statistical analyses, data wrangling, and visualizations were performed in the R environment (version 4.5.1). Single-cell transcriptomic analyses, including quality control, integration, dimensionality reduction, and clustering, were conducted using the Seurat R package (version 5.5.0) and its underlying matrix operations handled by the Matrix package (version 1.7.5). To mitigate the confounding effects of multiplets, doublet detection was performed using scDblFinder (version 1.21.2), supported by the scran (version 1.36.0) and scater (version 1.36.0) frameworks. Gene Ontology functional enrichment analyses were executed using clusterProfiler (version 4.16.0) and DOSE (version 4.2.0), utilizing the org.Hs.eg.db annotation database (version 3.21.0). Data manipulation and formatting were facilitated by dplyr (version 1.2.1), tidyr (version 1.3.1), plyr (version 1.8.9), reshape2 (version 1.4.5), and openxlsx (version 4.2.8.1). All graphical visualizations were generated using ggplot2 (version 4.0.3), patchwork (version 1.3.2), scales (version 1.4.0), and viridis (version 0.6.5), with foundational support from BiocGenerics (version 0.54.1). Raw sequencing data were processed using the 10x Genomics Cell Ranger pipeline (version 10.0.0). All custom scripts utilized for data processing, integration, and visualization are available in the Supplementary Information and have been deposited in a public GitHub repository.

### Surgical procedures for sciatic nerve defect model and graft implantation

All animal procedures were performed in accordance with protocols approved by the Institutional Animal Care and Use Committee of the Chinese Institute for Brain Research, Beijing (CIBR-IACUC), under approval number CIBR-IACUC-218, and conformed to the NIH guidelines for the care and use of laboratory animals. Female NSG (NOD.Cg-Prkdcscid Il2rgtm1Wjl/SzJ) mice (6-8 weeks old) were used as recipients for graft implantation. Animals were fasted for 6 h prior to surgery with free access to water. Anesthesia was induced by intraperitoneal injection of 1.25% tribromoethanol (Avertin, Aibei bio) at a dose of 0.3 mL/10 g body weight. Anesthetic depth was confirmed by the absence of toe pinch reflex. Mice were positioned in left lateral recumbency, and the surgical area (left hindlimb, lateral thigh from hip to knee) was shaved using an electric clipper and disinfected twice with povidone-iodine. Sterile drapes were applied to maintain aseptic conditions.

A longitudinal skin incision of approximately 1.5 cm was made at the midpoint between the greater trochanter and the lateral epicondyle of the knee, along the intermuscular plane between the biceps femoris and the gluteus maximus. Subcutaneous fascia was sharply dissected, and the biceps femoris and vastus lateralis were bluntly separated to expose the sciatic nerve. The nerve was carefully freed from surrounding connective tissue, extending proximally to the sciatic notch and distally to the popliteal bifurcation, exposing a total length of approximately 1.2 cm of the sciatic nerve, while preserving the adjacent vasculature.

A 7-mm segment of the sciatic nerve was sharply resected at the mid-thigh level using micro-scissors, creating an irreversible gap. The corresponding repair material, either autologous nerve (reversed and sutured), hollow silicone tube, or the Peri-nervoid graft, was placed between the proximal and distal nerve stumps. Under a 10× surgical microscope (Bondent, #DOM3000), the nerve stumps were coapted to the graft using 10-0 nylon sutures with epineurial sutures (four interrupted sutures per end), ensuring close apposition without torsion or tension.

The wound was irrigated with sterile saline to remove blood clots and debris. The muscle layer was closed with 4-0 absorbable sutures in a continuous pattern, and the skin was closed with 5-0 nylon interrupted sutures. The incision was disinfected with povidone-iodine after closure.

Postoperatively, mice were placed on a warming pad (37°C) for recovery and housed individually after full awakening. Meloxicam (4 mg/kg) was administered subcutaneously within 48 h post-surgery for analgesia. Incisions were disinfected daily with povidone-iodine until complete healing. Body weight, wound condition, and autotomy behavior were monitored daily for the first week post-surgery.

### Gait analysis

The gait of experimental animals was analyzed 70 DPT using the Catwalk automated gait analysis system, capturing footprints of the animals (n = 5 per group). The sciatic functional index (SFI) was calculated based on the animal footprints during walking using the Catwalk XT 10.6 software from Noldus. Three parameters of the footprints were evaluated for both the injured (E) and normal (N) sides, including toe spread (TS), intermediate toe spread (ITS), and print length (PL). The SFI was calculated using the Bain formula^31^: SFI = 109.5(ETS-NTS)/NTS -38.3(EPL-NPL)/NPL +13.3(EIT - NIT)/NIT - 8.8.

### Von Frey test

Mechanical allodynia was assessed using the von Frey filament test at 10 weeks post-transplantation (70 DPT) as previously described^32^. Mice (n = 5 per group) were individually placed on a wire-mesh grid floor in plastic cages and acclimated for 1 h. Calibrated von Frey filaments (TACTILE TEST AESTHESIO; Muromachi Kikai, Tokyo, Japan) with bending forces of 0.16, 0.4, 0.6, 2, 4, 8g were applied perpendicularly to the mid-plantar surface of the hind paw through the mesh floor for 3–5 s with slight buckling. Stimuli were presented in ascending or descending order according to the up-down method. In the absence of a withdrawal response, a stronger filament was applied; in the presence of withdrawal, a weaker filament was chosen. After the response threshold was first crossed (i.e., two responses straddling the threshold), four additional stimuli were applied to determine the 50% paw withdrawal threshold.

### Hot and cold plate tests

Thermal nociception was assessed using hot and cold plate tests at 70 DPT. For the hot plate test, mice were individually placed on a hot plate apparatus (Model 35100, Ugo Basile, Italy) enclosed by a transparent Plexiglas cylinder (20 cm diameter, 30 cm height), with the plate maintained at 55.0 ± 0.5°C. The latency to the first nociceptive response (hind paw licking, shaking, or flicking) was recorded, with a cut-off time of 60 s to prevent tissue damage. For the cold plate test, the same apparatus was used with the plate temperature maintained at 4.0 ± 0.5°C. The latency to the first clear hind paw lift (sustained for >1 s) or licking was recorded, with a cut-off time of 60 s. The plate surface was cleaned with 70% ethanol and dried between each animal.

### Electrophysiological analysis

At 70 DPT, electrophysiological assessment was performed to evaluate nerve conduction recovery. Mice were weighed and anesthetized by intraperitoneal injection of 1.25% tribromoethanol (Avertin) at a dose of 0.3 mL/10 g body weight (n = 5). The right sciatic nerve graft and the intact contralateral left sciatic nerve were carefully exposed through a complete dissection. A stimulating electrode was placed proximally on the graft, while recording electrodes were positioned in the target gastrocnemius muscle. A single stimulus of 3 mA was delivered, and the compound muscle action potential (CMAP) was recorded. The same parameters were applied to the contralateral (intact) side for normalization. CMAP latency and amplitude ratio (ipsilateral/contralateral) were analyzed for each subject.

### Histopathology and hematological analysis

To evaluate the in vivo biosafety of the implanted grafts, a comprehensive post-mortem analysis was performed at 70 DPT (n = 5). Whole blood was collected via cardiac puncture immediately after euthanasia. A portion of the blood was transferred to EDTA-coated tubes for complete blood count (CBC) analysis, and the remaining blood was centrifuged at 1,500 × g for 15 min at 4°C to obtain serum for serum biochemistry panel analysis, including liver and kidney function markers.

Major organs, including the heart, liver, spleen, lungs, and kidneys, were harvested and weighed. All tissue collection procedures were performed collaboratively by two investigators to ensure consistent processing times across organs and to minimize tissue degradation. Organs were fixed in 4% paraformaldehyde (PFA) for 24 h at 4°C, embedded in paraffin, sectioned, and stained with hematoxylin and eosin (H&E). Stained sections were examined under a light microscope to assess for the presence of inflammatory infiltration, necrosis, tumor formation, or any other histopathological abnormalities.

### Assessment of target muscle recovery

At 70 DPT, bilateral gastrocnemius muscles were excised (n = 5). The wet weight ratio was calculated as the mass of the injured side divided by that of the contralateral side. Muscles were fixed, processed, sectioned, and stained using a Masson staining kit (G1006, Servicebio) according to the manufacturer’s instructions.

### Transmission Electron Microscopy (TEM)

Samples were fixed with TEM fixative (Servicebio, #G1102) for 15 min at room temperature. After transfer to a dish with fresh fixative, the axon bundles were transversely cut using a scalpel, and tissue blocks were collected into EP tubes and processed at 4°C. The fixative was replaced with 1% OsO₄ (Ted Pella, #18456) and incubated at room temperature for 2 h with rotation. Samples were then stained with 1% uranyl acetate for 2 h with rotation, dehydrated through graded ethanol series (50%, 70% overnight at 4°C, 95%, and 100%), followed by two 15-min changes of acetone.

Samples were infiltrated with EMBed 812 (SPI, #90529-77-4) in acetone gradients (1:1 for 2–4 h at 37°C; 1:2 overnight at 37°C; pure resin for 5–8h at 37°C), embedded in pure resin, and polymerized at 60°C for >48 h. Semi-thin sections (1.5μm) were cut, stained with toluidine blue, and examined by light microscopy for region selection. Ultrathin sections (60–80nm) were collected onto formvar-coated copper grids, stained with 2% uranyl acetate in saturated alcohol for 8min (dark), rinsed in 70% ethanol and ultrapure water, followed by 2.6% lead citrate for 8min. After rinsing and air-drying overnight, grids were examined using a Hitachi HT7800 TEM.

### Cryopreservation and immunohistochemistry

Samples were fixed in 4% PFA for 12 h at room temperature, washed three times with PBS, and cryoprotected in 30% sucrose in PBS at 4°C until saturated. Samples were then embedded in OCT compound (Surgipath FSC 22 Blue, Leica, #3801481) and frozen. Cryosections (16 μm) were cut using a cryostat (Leica), washed with PBS to remove OCT, blocked with 10% normal goat serum (NGS) and 0.3% Triton X-100 in PBS for 1 h at room temperature, and incubated with primary antibodies in blocking solution overnight at 4°C. After washing, sections were incubated with Alexa Fluor-conjugated secondary antibodies (1:1,000; Life Technologies) for 2 h at room temperature, followed by DAPI nuclear staining. Primary antibodies used in this study are listed in Table 3.

### Quantitative real time PCR

For qPCR analysis, 5-8 spheroids were pooled per sample. mRNA was extracted using the RNeasy Plus Micro kit (QIAGEN, #74034), and cDNA was synthesized by reverse transcription with the HiScript III All-in-one RT SuperMix Perfect for qPCR (Vazyme, #R333-01). qPCR was then performed using the Taq Pro Universal SYBR qPCR Master Mix (Vazyme, #Q712-02) on a QuantStudio 3 system (Applied Biosystems). The primers used are listed in Table 4.

### Imaging and analysis

Bright-field images of cells and organoids were captured at various time points using an inverted phase-contrast microscope (Olympus). Stained slides were imaged with a confocal panoramic scanner (Pannoramic, 3DHISTECH). For tissue clearing, samples were embedded in 2% low-melting-point agarose in PBS, dehydrated sequentially through 50%, 70%, and 100% ethanol (twice), delipidated in pure dichloromethane (DCM), and finally incubated in dibenzyl ether (DBE) solution for refractive index matching. Cleared samples were immersed in a DBE-filled chamber and imaged on a Zeiss Lightsheet 7 microscope equipped with dual-sided 5×/0.1 NA illumination optics and a 5×/0.16 NA detection objective, using continuous light-sheet scanning with sequential single-side illumination. Image stacks were reconstructed with Zen Blue (Zeiss), and 3D volume rendering and analysis were performed using Imaris (version 10.2).

### Statistical analyses

Statistical analyses were performed using GraphPad Prism 10.1.2. Data are presented as mean ± SD. Comparisons among multiple groups were conducted using one-way analysis of variance (ANOVA) followed by Tukey’s multiple-comparison test. For comparisons between two groups, two-tailed unpaired Student’s t-tests were used. A *P* value < 0.05 was considered statistically significant.

## Acknowledgements

We thank Dr. Xiangyu Meng, Ms. Shan Zhang and Ms. Wenjing Wu (the Central Laboratory of Beijing Jishuitan Hospital, Capital Medical University) for assistance with cell process, confocal microscopy and flow cytometry; Together we thank Ms. Dan Zhang (Core Facility, Center of Biomedical Analysis, Tsinghua University) for technical support with whole mount immunolabeling, tissue clearing, lightsheet microscopy, data analysis and processing.

## Declaration of generative AI and AI-assisted technologies in the writing process

During the preparation of this work the authors used ChatGPT (OpenAI) to enhance the clarity, grammar, and readability of the text. After using this tool/service, the authors reviewed and edited the content as needed and take full responsibility for the content of the publication.

## Author contributions

Y.G. designed experiments, performed studies, analyzed data, and wrote the manuscript. R.H. performed studies, analyzed data, designed schematic illustrations, and wrote the manuscript. Z.X. conducted the differentiation and characterization of PSC-derived spinal cord organoids and performed animal experiments. Y.O. conducted the differentiation and characterization of PSC-derived dorsal root ganglion organoids, and analyzed sc-RNA sequencing data. H.L. performed hPSC-SCs differentiation, analyzed data. S.Z. conducted Von Frey assessments, hot and cold plate tests. X.L. conducted calcium imaging, and analyzed data. C.L. conducted histological staining.

Z.W. conceived the study idea, supervised data analysis and revised the manuscript. Y.W. conceived the study idea and revised the manuscript. S.C. con-ceptualized this study, provided guidance on experimental design, supervised data analysis, and revised the manuscript.

## Supplementary material

Supplemental material for this article is available online.

## Reference

1. Aguayo, A.J., Kasarjian, J., Skamene, E., Kongshavn, P., and Bray, G.M. (1977). Myelination of mouse axons by Schwann cells transplanted from normal and abnormal human nerves. Nature 268, 753–755. 10.1038/268753a0.

2. Cattin, A.L., Burden, J.J., Van Emmenis, L., Mackenzie, F.E., Hoving, J.J., Garcia Calavia, N., Guo, Y., McLaughlin, M., Rosenberg, L.H., Quereda, V., et al. (2015). Macrophage-Induced Blood Vessels Guide Schwann Cell-Mediated Regeneration of Peripheral Nerves. Cell 162, 1127–1139. 10.1016/j.cell.2015.07.021.

3. Stassart, R.M., Gomez-Sanchez, J.A., and Lloyd, A.C. (2024). Schwann Cells as Orchestrators of Nerve Repair: Implications for Tissue Regeneration and Pathologies. Cold Spring Harb Perspect Biol 16. 10.1101/cshperspect.a041363.

4. Jessen, K.R., and Arthur-Farraj, P. (2019). Repair Schwann cell update: Adaptive reprogramming, EMT, and stemness in regenerating nerves. Glia 67, 421 – 437. 10.1002/glia.23532.

5. Ouyang, Y., Yu, M., Zhang, T., Cheng, H., Zuo, L., Liu, H., Guan, Y., Liu, A., Chen, J., He, R., et al. (2025). Single-cell transcriptomic landscape of sciatic nerve after transection injury. J Neuroinflammation 22, 205. 10.1186/s12974-025-03514-3.

6. Rotshenker, S. (2011). Wallerian degeneration: the innate-immune response to traumatic nerve injury. J Neuroinflammation 8, 109. 10.1186/1742-2094-8-109.

7. Conforti, L., Gilley, J., and Coleman, M.P. (2014). Wallerian degeneration: an emerging axon death pathway linking injury and disease. Nat Rev Neurosci 15, 394 – 409. 10.1038/nrn3680.

8. Lopes, B., Sousa, P., Alvites, R., Branquinho, M., Sousa, A.C., Mendonça, C., Atayde, L.M., Lujs, A.L., Varejão, A.S.P., and Maurjcio, A.C. (2022). Peripheral Nerve Injury Treatments and Advances: One Health Perspective. Int J Mol Sci 23. 10.3390/ijms23020918.

9. Gu, X., Ding, F., and Williams, D.F. (2014). Neural tissue engineering options for peripheral nerve regeneration. Biomaterials 35, 6143–6156. 10.1016/j.biomaterials.2014.04.064.

10. Levi, A.D., and Bunge, R.P. (1994). Studies of myelin formation after transplantation of human Schwann cells into the severe combined immunodeficient mouse. Exp Neurol 130, 41–52. 10.1006/exnr.1994.1183.

11. Mosahebi, A., Fuller, P., Wiberg, M., and Terenghi, G. (2002). Effect of allogeneic Schwann cell transplantation on peripheral nerve regeneration. Exp Neurol 173, 213 – 223. 10.1006/exnr.2001.7846.

12. Monje, P.V. (2020). The properties of human Schwann cells: Lessons from in vitro culture and transplantation studies. Glia 68, 797–810. 10.1002/glia.23793.

13. Yamanaka, S. (2012). Induced pluripotent stem cells: past, present, and future. Cell Stem Cell 10, 678–684. 10.1016/j.stem.2012.05.005.

14. Majd, H., Amin, S., Ghazizadeh, Z., Cesiulis, A., Arroyo, E., Lankford, K., Majd, A., Farahvashi, S., Chemel, A.K., Okoye, M., et al. (2023). Deriving Schwann cells from hPSCs enables disease modeling and drug discovery for diabetic peripheral neuropathy. Cell Stem Cell 30, 632–647.e610. 10.1016/j.stem.2023.04.006.

15. Moss, K.R., Mi, R., Kawaguchi, R., Ehmsen, J.T., Shi, Q., Vargas, P.I., Mukherjee-Clavin, B., Lee, G., and Höke, A. (2024). hESC- and hiPSC-derived Schwann cells are molecularly comparable and functionally equivalent. iScience 27, 109855. 10.1016/j.isci.2024.109855.

16. Jessen, K.R., Mirsky, R., and Lloyd, A.C. (2015). Schwann Cells: Development and Role in Nerve Repair. Cold Spring Harb Perspect Biol 7, a020487. 10.1101/cshperspect.a020487.

17. Salzer, J., Feltri, M.L., and Jacob, C. (2024). Schwann Cell Development and Myelination. Cold Spring Harb Perspect Biol 16. 10.1101/cshperspect.a041360.

18. Panzer, K.V., Burrell, J.C., Helm, K.V.T., Purvis, E.M., Zhang, Q., Le, A.D., O’Donnell, J.C., and Cullen, D.K. (2020). Tissue Engineered Bands of Büngner for Accelerated Motor and Sensory Axonal Outgrowth. Front Bioeng Biotechnol 8, 580654. 10.3389/fbioe.2020.580654.

19. Andersen, J., Revah, O., Miura, Y., Thom, N., Amin, N.D., Kelley, K.W., Singh, M., Chen, X., Thete, M.V., Walczak, E.M., et al. (2020). Generation of Functional Human 3D Cortico-Motor Assembloids. Cell 183, 1913–1929.e1926. 10.1016/j.cell.2020.11.017.

20. Mazzara, P.G., Muggeo, S., Luoni, M., Massimino, L., Zaghi, M., Valverde, P.T., Brusco, S., Marzi, M.J., Palma, C., Colasante, G., et al. (2020). Frataxin gene editing rescues Friedreich’s ataxia pathology in dorsal root ganglia organoid-derived sensory neurons. Nat Commun 11, 4178. 10.1038/s41467-020-17954-3.

21. Hagemann, C., Moreno Gonzalez, C., Guetta, L., Tyzack, G., Chiappini, C., Legati, A., Patani, R., and Serio, A. (2022). Axonal Length Determines Distinct Homeostatic Phenotypes in Human iPSC Derived Motor Neurons on a Bioengineered Platform. Adv Healthc Mater 11, e2101817. 10.1002/adhm.202101817.

22. Guan, Y., Jia, Z., Xiong, X., He, R., Ouyang, Y., Liu, H., Liang, L., Meng, X., Zhang, R., Guan, C., et al. (2025). Tissue-specific extracellular matrix for the larger-scaled expansion of spinal cord organoids. Mater Today Bio 31, 101561. 10.1016/j.mtbio.2025.101561.

23. Huang, Z., Powell, R., Kankowski, S., Phillips, J.B., and Haastert-Talini, K. (2023). Culture Conditions for Human Induced Pluripotent Stem Cell-Derived Schwann Cells: A Two-Centre Study. Int J Mol Sci 24. 10.3390/ijms24065366.

24. Ben-David, U., and Benvenisty, N. (2011). The tumorigenicity of human embryonic and induced pluripotent stem cells. Nat Rev Cancer 11, 268–277. 10.1038/nrc3034.

25. Dombrowski, Y., O’Hagan, T., Dittmer, M., Penalva, R., Mayoral, S.R., Bankhead, P., Fleville, S., Eleftheriadis, G., Zhao, C., Naughton, M., et al. (2017). Regulatory T cells promote myelin regeneration in the central nervous system. Nat Neurosci 20, 674–680. 10.1038/nn.4528.

26. Deuse, T., Hu, X., Gravina, A., Wang, D., Tediashvili, G., De, C., Thayer, W.O., Wahl, A., Garcia, J.V., Reichenspurner, H., et al. (2019). Hypoimmunogenic derivatives of induced pluripotent stem cells evade immune rejection in fully immunocompetent allogeneic recipients. Nat Biotechnol 37, 252–258. 10.1038/s41587-019-0016-3.

27. Kim, H.S., Lee, J., Lee, D.Y., Kim, Y.D., Kim, J.Y., Lim, H.J., Lim, S., and Cho, Y.S. (2017). Schwann Cell Precursors from Human Pluripotent Stem Cells as a Potential Therapeutic Target for Myelin Repair. Stem Cell Reports 8, 1714–1726. 10.1016/j.stemcr.2017.04.011.

28. Chambers, S.M., Qi, Y., Mica, Y., Lee, G., Zhang, X.J., Niu, L., Bilsland, J., Cao, L., Stevens, E., Whiting, P., et al. (2012). Combined small-molecule inhibition accelerates developmental timing and converts human pluripotent stem cells into nociceptors. Nat Biotechnol 30, 715–720. 10.1038/nbt.2249.

29. Schwartzentruber, J., Foskolou, S., Kilpinen, H., Rodrigues, J., Alasoo, K., Knights, A.J., Patel, M., Goncalves, A., Ferreira, R., Benn, C.L., et al. (2018). Molecular and functional variation in iPSC-derived sensory neurons. Nat Genet 50, 54–61. 10.1038/s41588-017-0005-8.

30. Mukherjee-Clavin, B., Mi, R., Kern, B., Choi, I.Y., Lim, H., Oh, Y., Lannon, B., Kim, K.J., Bell, S., Hur, J.K., et al. (2019). Comparison of three congruent patient-specific cell types for the modelling of a human genetic Schwann-cell disorder. Nat Biomed Eng 3, 571 – 582. 10.1038/s41551-019-0381-8.

31. Bain, J.R., Mackinnon, S.E., and Hunter, D.A. (1989). Functional evaluation of complete sciatic, peroneal, and posterior tibial nerve lesions in the rat. Plast Reconstr Surg 83, 129–138. 10.1097/00006534-198901000-00024.

32. Liu, H., Ouyang, Y., Wang, B., Zhang, X., Guan, Y., He, R., Cui, Y., Wang, J., Yao, Q., Tan, Y., et al. (2026). Mesenchymal Stem Cell-Derived Apoptotic Micro-Vesicles Repaired Sciatic Nerve Defect by Regulating Early Inflammatory Microenvironment and Promoting Angiogenesis. Adv Healthc Mater 15, e04087. 10.1002/adhm.202504087.

