## supplementary information for "Generation and application of human pluripotent stem cells derived peri-nervoids"

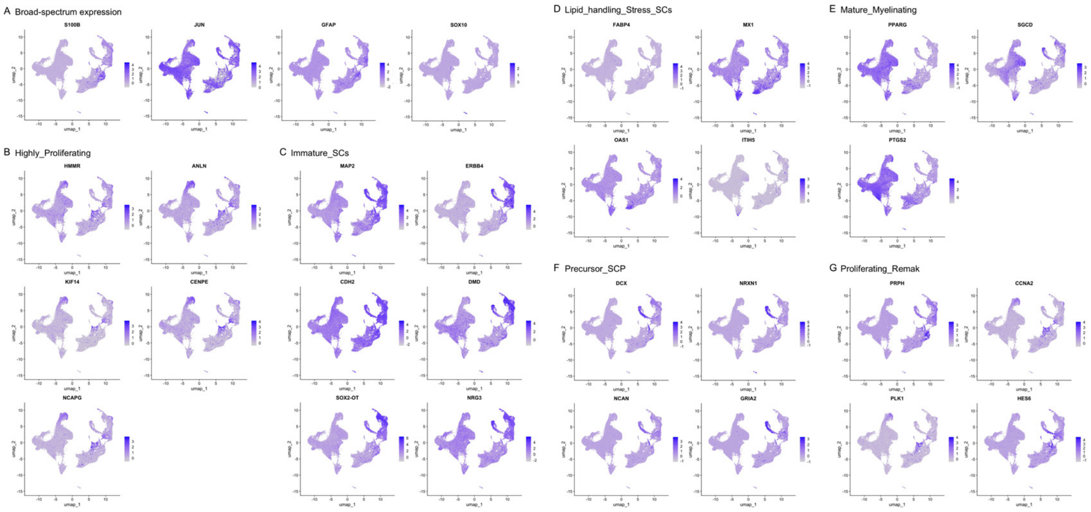


**Supplementary Fig. 1. Enrichment of specific marker genes in different Schwann cell subtypes.**

**A.** Distribution of typical markers genes in Schwann cells.

**B-G.** Feature plots illustrating the expression of each subtype of Schwann cell specific marker genes.

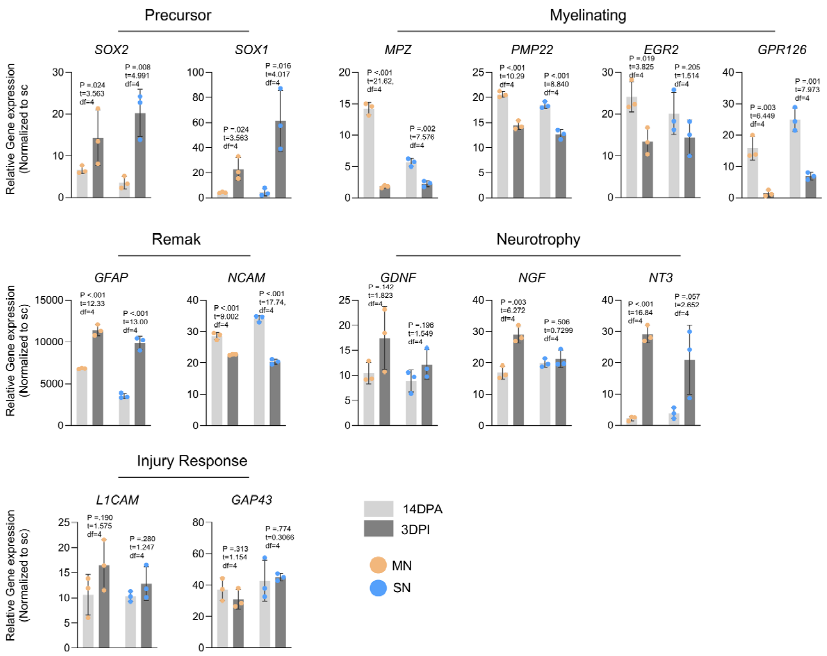


**Supplementary Fig. 2. qPCR validation of SCs transcriptional dynamics in response to axonal co-culture and injury.** Compared with SCs cultured alone (sc), all examined genes were upregulated in both MN-PNOs and SN-PNOs at 14 DPA and 3 DPI. At 3 DPI vs. 14 DPA, precursor markers (*SOX1*, *SOX2*) and neurotrophic factors (*GDNF*, *NGF*, *NT3*) were further upregulated in both MN and SN groups, whereas myelination markers (*MPZ*, *PMP22*, *EGR2, GPR126*) were decreased. Data are shown as mean ± SD. (n = 3 biological replicates per time point, two-sided unpaired Student's *t* test).

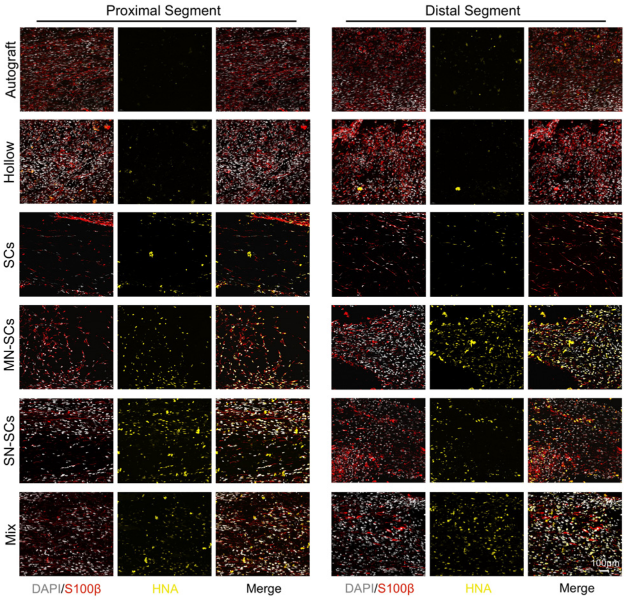


**Supplementary Fig. 3. Transplanted human Schwann cells survive and distribute in both proximal and distal segments of PNO grafts at 14 DPT.**

Representative images of proximal (left) and distal (right) graft segments show distribution and survival of transplanted human-derived Schwann cells ($HNA^+$ /$S100\beta^+$) within the regenerating nerve bridge.

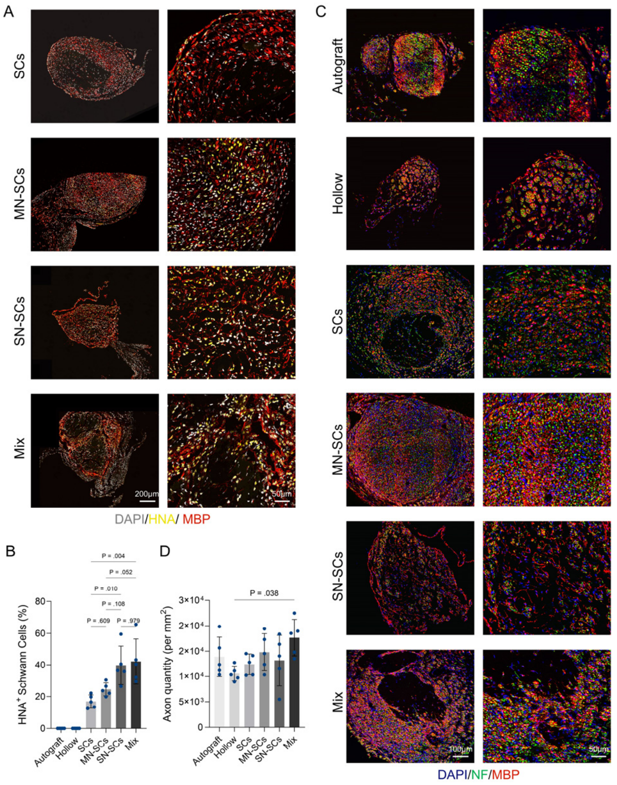


**Supplementary Fig. 4. Cross-sectional analysis confirms comparable human Schwann cell integration and axonal density among neuronal-containing PNO grafts at 70 DPT.**

**A.** Representative cross-sections stained for HNA (green) and MBP (red).

**B.** Quantification of $HNA^+$/$MBP^+$ human Schwann cell proportions (n = 5 animals each group, one-way ANOVA: F (3, 16) = 7.68; P = 0.002).

**C.** Representative cross-sections stained for NF (green) and MBP (red).

**D.** Quantification of axonal density (n = 5 animals each group, one-way ANOVA: F (5, 24) = 2.36; P = 0.071).

Data are shown as mean ± SD.

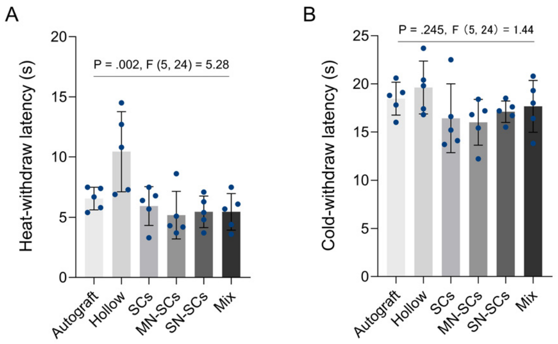


**Supplementary Fig. 5. Thermal sensory recovery is comparable among cell-containing groups and autografts.**

**A.** Quantitative analysis of heat withdrawal latency (n = 5 biological replicates, one-way ANOVA followed by Tukey's multiple-comparisons test: Hollow vs SCs: p = 0.013; Hollow vs MN-SCs: p = 0.003; Hollow vs SN-SCs: p = 0.005; Hollow vs MIX: p = 0.005. Autograft vs all cell-containing groups: n.s. (p > 0.05); all cell-containing groups vs each other: n.s. (p > 0.05)). Data are shown as mean ± SD. n.s., not significant.

**B.** Quantitative analysis of cold withdrawal latency (n = 5 biological replicates, one-way ANOVA followed by Tukey's multiple-comparisons test, no significant differences were detected among any of the experimental groups (all adjusted p > 0.99)). Data are shown as mean ± SD.

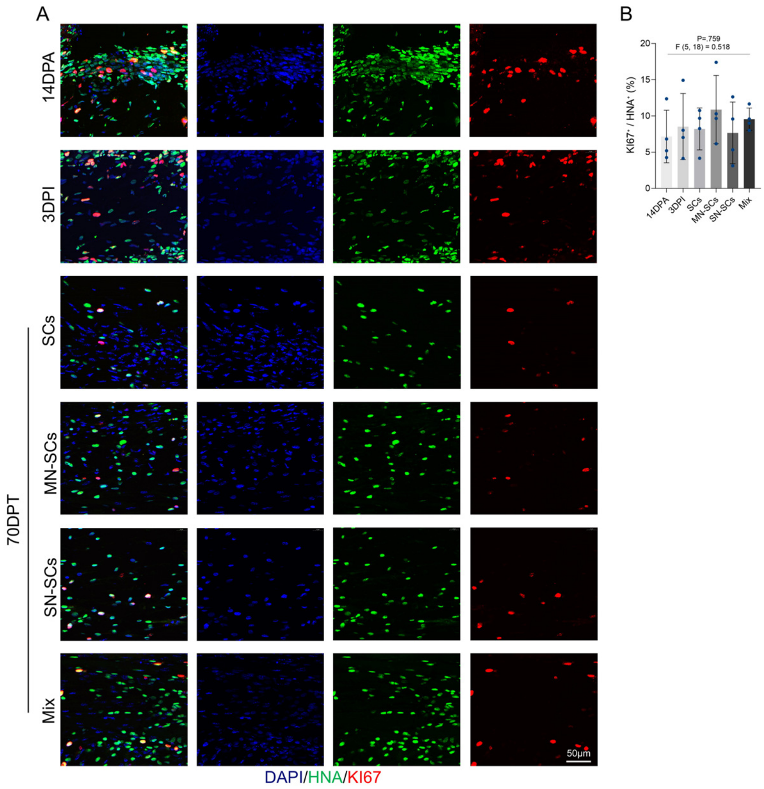


**Supplementary Fig. 6. Transplanted human Schwann cells do not undergo abnormal proliferation in vitro or in vivo.**

**A.** Immunofluorescence staining of PNOs and grafts for HNA (human nuclei, green) and KI67 (proliferation marker, red), with DAPI counterstain (blue).

**B.** The percentage of $HNA^+$ /$KI67^+$ double-positive cells were consistently low, with no significant differences between groups (one-way ANOVA), indicating that transplanted human Schwann cells did not undergo abnormal proliferation or neoplastic transformation (n = 4 biological replicates). Data are shown as mean ± SD.

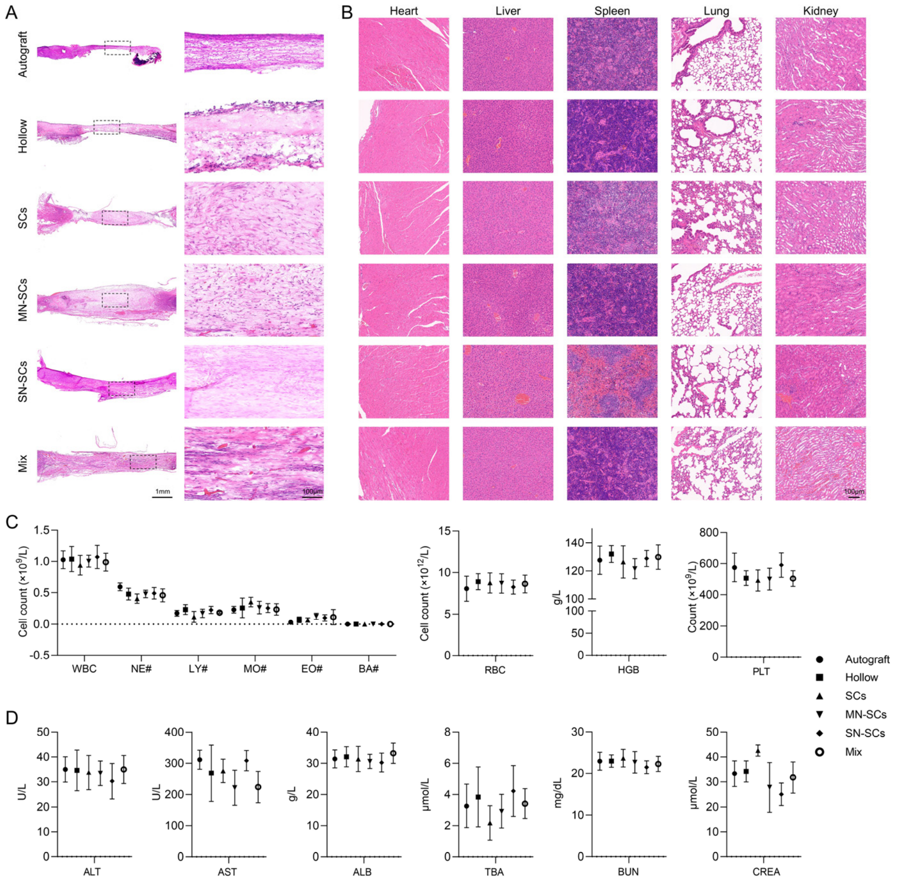


**Supplementary Fig. 7. Biosafety assessment confirms no systemic toxicity or organ damage following Peri-nervoid transplantation.**

**A.** Representative H&E staining of mid-graft sections at 70 DPT, showing normal tissue architecture with no evidence of tumor formation, hypercellularity, or atypical cell clusters.

**B.** H&E staining of major organs (heart, liver, spleen, lung, and kidney) from representative animals across all experimental groups, demonstrating no signs of inflammation, necrosis, or tumorigenesis.

**C.** Complete blood count analysis including white blood cells (WBC), neutrophils (NE#), lymphocytes (LY#), monocytes (MO#), eosinophils (EO#), basophils (BA#), red blood cells (RBC), hemoglobin (HGB), and platelets (PLT) (n = 5 biological replicates).

**D.** Serum biochemistry panel showing liver function markers (ALT, AST, ALB, TBA) and kidney function markers (BUN, CREA) (n = 5 biological replicates). No significant differences were detected among any of the groups for all hematological and biochemical parameters evaluated (one-way ANOVA).

Data are presented as mean ± SD.
